# High-throughput phenotyping reveals natural variation in tomato pollen responses to heat stress

**DOI:** 10.64898/2026.11.19.767983

**Authors:** Cedar Warman, Calvin Perkins, Sara McKinley, Berenice Gonzalez, Kekhrie Tsurho, Steven E. Smith, James Pease, Ravishankar Palanivelu

## Abstract

Scalable measurements of pollen function are needed to investigate genetic variation in reproductive responses to heat. We developed a high-throughput phenotyping framework integrating automated bright-field microscopy, object detection, individual-tube tracking, and temporal analysis. We quantified pollen performance at 26 and 34 °C across 218 tomato accessions and wild relatives by analyzing approximately 300,000 images collected over two-hour growth periods. Automated counts of pollen classes - aborted pollen grains, ungerminated, germinated, and burst pollen grains, and pollen tube tips - agreed closely with manual counts (*R*^2^ = 0.915–0.967). Heat stress shortened the mean time to 50% germination from 24.0 to 20.5 minutes but increased mean percent bursting at 2 hours from 14.9% to 83.1%. A burst integral metric combined the extent and timing of bursting and revealed apparently substantial variation among accessions. Mean pollen tube length was shorter at 34 °C than at 26 °C in 72% of accessions and was equal or greater at 34 °C in the remaining 28%. Individual-tube tracking revealed contrasting growth-speed distributions in representative thermosensitive and thermotolerant accessions, including persistence of faster-growing tubes in the thermotolerant Tamaulipas variety. This framework enables population-scale analysis of dynamic pollen traits and provides quantitative phenotypes for investigating the genetic basis of pollen thermotolerance.

## Introduction

Fertilization in flowering plants relies on tightly coordinated reproductive processes essential for seed and fruit formation, generating new plants, and ultimately for sustaining global crop yields (Amasino *et al*., 2017; Pabón-Mora *et al*., 2022). During the final stages of flowering plant reproduction, pollen grains are released from the anthers and deposited on the stigma, where they germinate and produce a pollen tube (Johnson *et al*., 2019). Pollen tubes grow through the stigmatic surface and extend into the pistil, where intrinsic and extrinsic molecular cues guide them toward the ovules through tip-directed growth (Adhikari *et al*., 2020). Upon reaching an ovule, a pollen tube delivers sperm cells, enabling fertilization and subsequent seed and fruit development.

The pollen tube cell wall must maintain a delicate balance between flexibility and integrity as this single cell often lengthens from micrometers to several millimeters in length. Upon reaching the female gametophyte, the pollen tube tip interacts with the synergid cells and exchanges cell-specific signaling ligands that determine reproductive compatibility (Kim *et al*., 2021). Successful pollen tube reception by the synergid cells triggers the rupture of the pollen tube tip, releasing its cytoplasm and sperm cells into the space between the egg and central cell. One sperm cell fuses with the egg cell, and the other fuses with the central cell, completing double fertilization. Thus, pollen tube rupture is an essential, tightly regulated step required for successful fertilization. Importantly, pollen tube rupture must occur at the correct time and location within the female gametophyte. Successful pollen tube germination, growth, and rupture reflect key cellular processes, including cell wall integrity, turgor regulation, and signaling-dependent pollen tube reception, which directly determine sperm delivery, fertilization, and seed formation. Defects in any aspect of pollen tube growth, including premature or delayed bursting, can compromise fertilization success in flowering plants and reduce seed set (Noble and Palanivelu, 2020).

Heat stress causes significant defects in pollen tube growth, making this phase one of the most temperature-sensitive stages of plant reproduction and leading to reduced seed set and yield across multiple crop species (Deryng *et al*., 2014; Hatfield and Prueger, 2015; Raza *et al*., 2019; Ouonkap *et al*., 2024). Global temperatures have reached record highs and are projected to continue rising, prompting concerns about the impact of more frequent and prolonged heat stress on crop yield (Zhao *et al*., 2017; Voosen, 2021). Pollen and pollen tubes are often regarded as the most heat-sensitive components of plant reproduction (Zinn et al., 2010). This vulnerability is attributed to their developmental cycle of dehydration and subsequent rehydration, and the highly dynamic growth of pollen tube tips, which demands rapid and precise cytosolic, membrane, and cell wall dynamics that are challenging to maintain under heat stress (Sze et al., 2021, 2024; Perkins et al., 2025). Recent studies have examined pollen tube fitness under heat stress in different flowering plants, revealing a range of heat-sensitive phenotypes, including reduced pollen viability, grain number, germination, and tube length, as well as increased pollen tube bursting (Saini *et al*., 1983; Hedhly *et al*., 2009; Muhlemann *et al*., 2018; Begcy *et al*., 2019; Chaturvedi *et al*., 2021; Li *et al*., 2024).

*Solanum lycopersicum* (tomato) is emerging as an ideal model for investigating pollen tube growth under heat stress, owing to its economic importance, suitability for precision breeding, and diverse germplasm shaped by natural and artificial selection (Muhlemann *et al*., 2018; Rutley *et al*., 2021; Postiglione *et al*., 2024; Ouonkap *et al*., 2024). Throughout domestication, farmers and breeders have selected tomatoes for a wide array of traits, including heat tolerance, defined as the ability to set seed and fruit under high temperatures. Comparison of heat-tolerant and heat-sensitive tomato cultivars shows that reduced seed set at high temperatures is directly linked to impaired pollen tube growth (Zhou *et al*., 2017; Rutley *et al*., 2021; Ouonkap *et al*., 2024). *In vivo*, heat-sensitive pollen tubes fail to reach the ovary, reducing fertilization and seed formation. *In vitro*, heat-sensitive pollen also shows reduced germination and increased grain and tube bursting at high temperatures compared with heat-tolerant counterparts. Tolerance to heat stress in pollen tubes has been attributed to the regulation of cytosolic oxidation state and cell wall integrity (Muhlemann *et al*., 2018; Ouonkap *et al*., 2024; Postiglione *et al*., 2024; Almasaud *et al*., 2026), but specific genetic determinants underlying pollen heat tolerance remain largely unknown. Despite extensive characterization of heat-sensitive phenotypes, the relationship between pollen tube cellular behaviors under heat stress and the underlying genetic variation remains poorly resolved, largely because scalable, quantitative phenotyping approaches are lacking.

Genomic analyses and genome editing offer efficient tools to engineer heat tolerance into sensitive crop varieties while maintaining favorable agronomic traits, enabling targeted improvement in crop resilience. Cultivated tomatoes and their wild relatives have adapted to diverse environments, accumulating a wide range of traits through evolution, domestication, and breeding. This phenotypic diversity provides a valuable resource for identifying genomic variants that underlie traits of interest. Variation in many critical reproductive and physiological traits often results from multiple causal genomic variants. Power to detect these causal variants typically increases with sample size, underscoring the importance of surveying many individuals (Hong and Park, 2012). The extensive genetic resources available in tomato can be used to identify causal variants underlying quantitative traits through population-scale genomic analyses (Fernandez-Pozo *et al*., 2015). However, screening more individuals creates a major bottleneck in generating robust phenotypic datasets, underscoring the need for high-throughput phenotyping methods.

Pollen tube growth offers a promising phenotype for high-throughput screening. I*n vitro* assays that isolate the male gametophyte from surrounding floral tissues can assess pollen tube growth and its direct fitness effects, allowing observation of dynamic growth under a microscope (Palanivelu and Johnson, 2010). While these assays do not account for the biological context of floral tissues, they offer a useful model for rapidly screening pollen fitness under controlled conditions. *In vitro* pollen tube growth can be completed within hours in liquid media, making it suitable for integration into high-throughput, automated imaging systems that enable rapid, large-scale data collection. Historically, researchers have quantified *in vitro* pollen growth assays manually using GUI-based image analysis software, measuring traits such as pollen germination, tube growth, and tube bursting by hand (Schindelin *et al*., 2012). Manual scoring is practical for small datasets and endpoint measurements, but the effort increases with each additional image and time point. Deep learning offers a promising solution, as *in vitro* pollen assay datasets present an ideal opportunity for automation and enable faster, more scalable trait quantification through neural network training (Warman and Fowler, 2021).

Recent advances in computer vision have enabled automated measurements of pollen traits, ranging from grain shape and size to functional traits such as germination and tube growth. Deep learning–based tools can classify pollen grains across taxa with high accuracy using morphology in microscopy images, providing a scalable alternative to manual identification by specialists (Yu *et al*., 2024). Building on these advances, researchers have widely applied image analysis to assess pollen viability, particularly through staining-based methods. Automated microscopy techniques using viability dyes such as fluorescein diacetate and propidium iodide have enabled classification of viable, dead, and sterile pollen with high throughput and statistical power (Ascari *et al*., 2020), and deep learning frameworks such as PollenDetect have further accelerated viability assessment with high accuracy and efficiency (Tan *et al*., 2022). Staining assays for pollen viability have recently been combined with object detection models to enable field-level screening, facilitating the identification of genes associated with superior maize pollen viability at elevated temperatures (Li et al., 2026). (Li *et al*., 2026).

In parallel, pollen viability has also been evaluated through *in vitro* germination assays that provide a more functional assessment of pollen performance. Deep learning approaches have been successfully applied to quantify pollen germination frequency and pollen tube length directly from microscopy images (Zhang *et al*., 2023), and machine learning methods have been developed to estimate germination rates across experimental conditions in species such as *Capsicum annuum* (Yamazaki *et al*., 2023). Güçlü (2025) measured both pollen germination and tube length *in vitro* across a range of assay conditions and used deep learning to predict germination rates from incubation temperature, duration, and germination medium. More recently, semi-automated tools such as TubeTracker have enabled measurement of multiple pollen performance parameters, including germination timing, tube elongation rate, and integrity, using time-lapse imaging, advancing comprehensive analysis of pollen behavior (Ouonkap *et al*., 2025). However, these studies have generally been limited to a small number of genotypes, typically under one or two environmental conditions, and have focused on a restricted set of phenotypic parameters.

Despite these advances, most existing approaches focus on a limited subset of pollen traits, often at a single time point or within a small number of genotypes, and cannot integrate multiple phenotypic dimensions across large, genetically diverse sets of accessions. In particular, current methods rarely capture the full temporal progression of pollen germination, growth, and failure, nor do they enable population-scale comparisons under environmental stress conditions. As a result, our understanding of how dynamic pollen behaviors vary across natural diversity and contribute to stress tolerance remains limited. Here, we address these gaps by developing a high-throughput, computer vision–based phenotyping framework that integrates object detection, tracking, and time-resolved analysis to quantify multiple pollen performance traits simultaneously across a diverse sample of tomato and wild relatives under control and heat-stress conditions.

This framework enables quantitative dissection of pollen tube growth dynamics at a scale that is otherwise difficult to attain, linking pollen phenotypes to the genetic variation underlying reproductive thermotolerance. Object detection models combine classification and localization tasks using convolutional neural networks (CNNs), making them well suited to this type of image data (Lamichhane *et al*., 2025). By applying transfer learning to an object detection architecture, we show that deep learning enables automated, high-throughput quantification of pollen traits with strong accuracy and scalability. This framework reduces the labor and subjectivity of manual analysis and provides a reproducible method for evaluating reproductive response to heat stress across genetically diverse germplasm.

Using this approach, we identify substantial natural variation in pollen thermotolerance and show that heat stress differentially impacts key aspects of pollen tube biology, including germination dynamics, tube growth, and pollen tube integrity *in vitro*. In particular, we demonstrate that heat stress accelerates pollen germination, increases pollen tube rupture, and alters growth in ways that may differ among accessions. By introducing integrative metrics that capture both the extent and temporal dynamics of pollen tube failure, we provide an additional quantitative framework for comparing thermotolerance across genotypes. The resulting phenotypic data are robust and enable downstream genotype-phenotype analyses, including identifying genetic targets to improve crop thermotolerance and advance agriculture through genome editing and precision breeding.

## Materials and Methods

### Diversity panel and germplasm collection

We assembled a genetically diverse set (panel) of 218 tomato accessions primarily from two sources: the Varitome project (Gao et al., 2019; Mata-Nicolás et al., 2020) and a collection of accessions sequenced using long-read technology (Alonge et al., 2020) (Supplemental Table S1). We also included additional tomato accessions known to be thermotolerant but not in these two datasets (Supplemental Table S1). The panel consisted of *Solanum lycopersicum* (171 accessions), *S. pimpinellifolium* (39 accessions), *S. cheesmaniae* (4 accessions), and *S. galapagense* (4 accessions). The code used to generate a map of accessions is available here: https://github.com/cedarwarman/tomato_pollen_phenotyping/blob/main/R/map_accessions.R. Tomato germplasm was obtained from the C.M. Rick Tomato Genetics Resource Center (https://tgrc.ucdavis.edu/), the USDA National Plant Germplasm System (https://www.ars-grin.gov/), and the laboratories of Dr. Esther van der Knaap (University of Georgia) and Dr. Zachary Lippman (Cold Spring Harbor Laboratory).

### Genomic data quality control

Because the diversity panel combined publicly available data from multiple projects and two sequencing platforms, we preprocessed the raw reads to minimize technology-specific biases and ensure consistent data quality before variant calling across accessions. Raw FASTQ files were downloaded from the NCBI SRA database using the nf-core pipeline, fetchngs (v1.13.0dev, commit 3ace2a6; Patel *et al*., 2021). The diversity panel included a mix of accessions sequenced with Illumina short-read (146 samples) and Oxford Nanopore long-read (63 samples) technologies. To remove systemic biases from differences in sequencing technology, we split Oxford Nanopore long-read samples into 150bp fragments using the sliding window function in the sequence processing toolkit, seqkit (v2.13.0; Shen *et al*., 2024; Schiffer *et al*., 2025). We trimmed adapter sequences from the Illumina short reads with the preprocessing tool fastp (v0.22.0; Chen *et al*., 2018). We aligned both fragmented long reads and trimmed short reads to the SL4.0 reference genome (Sol Genomics Network) using bwa-mem2 (v2.2.1; Vasimuddin *et al*., 2019). We then sorted and indexed the output alignments and checked alignment quality with samtools (v1.19; Li *et al*., 2009). Quality control metrics related to genomic data processing are available in Supplemental Table S2. All scripts relevant for reproducing the genomic analyses are available at https://github.com/CJPerkins1/pollen_phenotyping_paper_popgen_methods.git

### Variant calling

To generate a consistent Single Nucleotide Polymorphism (SNP) set for analyses across sequencing platforms, we called variants separately in the short-and long-read datasets and retained SNPs detected in both datasets. SNPs were called from the aligned reads using bcftools mpileup/call version 1.19 with settings set to: maximum depth of 100, minimum mapping quality of 25 for short reads and 20 for long reads, minimum base quality of 20, the multiallelic model, and variant-only positions (Danecek *et al*., 2021). Because indel rates differ between short-and long-read sequencing technologies, we called only SNPs. We normalized output vcf files with bcftools norm and used the normalized files to identify intersecting loci between short-and long-read datasets. Intersecting SNPs between datasets were identified with the *intersect* tool from *bcftools* and then merged. A total of 10,228,773 intersecting SNPs were used for downstream analyses. For population genomic analyses, we filtered the genome-wide SNPs with bcftools for quality, missing data, minor allele frequency, and depth.

### Population genomic analyses

To characterize population structure within the diversity panel, we combined metric multidimensional scaling (MDS) of pairwise genetic distances with model-based ancestry estimation using ADMIXTURE. This analysis is essential for generating a population context to interpret variation in pollen thermotolerance, including differences associated with shared ancestry or geographic origin. A genome-wide identity-by-state (IBS) distance matrix was calculated in PLINK v1.90b6.19 using the quality-controlled SNP set (3,122,743 variants, 208 individuals; total genotyping rate = 0.984) with the -- distance square 1-ibs command, producing a symmetric N × N matrix of pairwise genetic distances (1 − IBS) between all individuals. MDS was performed on this matrix using the SMACOF algorithm implemented in scikit-learn v1.6.1 (sklearn.manifold.MDS), fitting a two-dimensional solution from 10 random initializations (maximum 10,000 iterations, convergence tolerance 1 × 10⁻⁹, random seed 42) and retaining the solution with the lowest stress (Kruskal’s stress-1 = 0.163). Because MDS solutions are invariant to rotation and reflection, we rotated the final configuration onto the first two classical MDS axes using orthogonal Procrustes analysis to provide a consistent orientation; this does not alter inter-sample distances. We colored accessions by genotype group (Supplemental Table S1), combining all SLL cultivar subgroups (vintage, heirloom, fresh-market, processing, and modern) as SLL (modern).

We plotted MDS coordinates with a custom Python script. We ran ADMIXTURE v1.3.0 (Alexander *et al*., 2009) on the pruned dataset for *K* = 1 through 10 using 5-fold cross-validation. Cross-validation error decreased across the tested range (*K* = 1: 0.748 to K = 10: 0.442) without reaching a global minimum; however, the rate of decrease slowed substantially beyond *K* = 4, indicating diminishing returns from additional clusters. We therefore selected K = 4 as the most parsimonious model capturing the major population structure. We plotted ancestry proportions inferred by ADMIXTURE at *K* = 4 in order of dominant ancestry cluster.

### Plant growth conditions

Before germination, we sequentially surface-sterilized seeds with 18% hydrochloric acid (EMD Inc.) for 20 minutes, 10% (g/ml) trisodium phosphate (Savogran Company) for 20 minutes, and 2.7% liquid bleach (sodium hypochlorite, Clorox Company) for 20 minutes. Following sterilization, seeds were plated on damp paper towels in 100 x 15 mm Petri dishes (Midland Scientific), sealed with Parafilm (Bemis Company Inc.), and incubated in darkness at 28 °C until germination (2-3 days).

Germinated seedlings were transplanted into 2” pots (Greenhouse Megastore) containing a 4:1 (v/v) mixture of Pro-mix LP15 soil (Premier Tech Horticulture) and perlite (Vigoro Corporation). Plants were grown for approximately 2 weeks in growth chambers (Conviron) under a 13-h light/11-h dark photoperiod at 24 °C (day) and 17 °C (night), with light provided by VL-1 LED grow lights at approximately 500 µmol m^-2^ s^-1^ (Volt Lighting). Plants in 2” pots were not fertilized. Seedlings were transplanted into 3.5-inch pots (Greenhouse Megastore) using the same soil mixture and growth conditions. Plants in 3.5” pots were fertilized with 5-11-26 fertilizer (ICL Peters) at 100 ppm nitrogen, and elemental sulfur (Bonide Products LLC) was applied as needed to control powdery mildew. After approximately 4 weeks, seedlings were transplanted to 2-gallon pots (Greenhouse Megastore) filled with Pro-mix LP15 soil (Premier Tech Horticulture) and moved to a greenhouse (University of Arizona, Tucson, AZ). We fertilized greenhouse-grown plants with 5-11-26 fertilizer (ICL Peters) at 150 ppm nitrogen using a Dosatron D14MZ2 injector (Dosatron International) and maintained them until flowering.

### Plant imaging

To document morphological diversity and provide a reference for researchers working with this panel, we imaged all accessions at the seedling and flowering stages (Supplemental Figures 1A–B) and a subset at the fruiting stage (Supplemental Figure 1C). Original images of seedlings, flowers, and fruits used in the montages of Supplemental Figure 1 are available at https://figshare.com/s/84b864bbad133c780540 during review, and it will be available at https://doi.org/10.25422/azu.data.34845574 after the manuscript is published. We also shared the images and accession information with the C. M. Rick Tomato Genetics Resource Center at UC Davis for inclusion alongside the corresponding germplasm records if appropriate. We imaged seedlings approximately 5 weeks after germination using a Raspberry Pi 4 (Raspberry Pi Foundation) equipped with a Raspberry Pi HQ Camera (Raspberry Pi Foundation) and a 6mm wide-angle lens (Adafruit). We placed plants in a 40 cm x 40 cm light box (Finnhomy) for image acquisition. Images were captured with the “raspistill” utility and saved as PNGs with the “--encoding png” option. Radial lens distortion was corrected using the OpenCV Python library following the camera calibration and undistortion workflow described here: https://docs.opencv.org/4.5.3/dc/dbb/tutorial_py_calibration.html. The de-distortion code is available at: https://github.com/cedarwarman/sporophyte_phenotyping/. We captured images of flowers and fruits on greenhouse-grown plants using an iPhone 13 Pro (Apple Inc.). We obtained fruit images only for a subset of accessions that successfully set fruit.

### Pollen microscopy and image acquisition

#### Microscopy

We performed high-throughput pollen phenotyping using a Leica DMI6000B inverted confocal microscope integrated with the Leica SP5 system to enable automated image acquisition. We used the Leica Application Suite (LAS) to configure, control, and operate the microscope system. Original images of pollen used in this study are available at https://figshare.com/s/84b864bbad133c780540 during review, and it will be available at https://doi.org/10.25422/azu.data.34845574 after the manuscript is published.

We captured these pollen images using two detection systems. We acquired about two-thirds of the images using the Leica confocal detector (PMT Trans) (camera 1) and captured the remaining images using a Leica DFC340 FX digital camera (camera 2). Images were acquired using confocal settings with an Acousto-Optic Tunable Filter at 543 nm, 25% laser power, and a scan speed of 600 Hz.

#### Pollen growth media

Liquid pollen growth media (PGM) consisted of 24% (g/mL) polyethylene glycol 4000 (CAS#25322-68-3), 0.01% (g/mL) boric acid (CAS#10043-35-3), 2% sucrose (CAS#57-50-1), 20mM HEPES, pH 6 (CAS#7365-45-9), 3mM calcium nitrate tetrahydrate (CAS#13477-34-4), 0.02% (g/mL) magnesium sulfate heptahydrate (CAS#10034-99-8), 0.01% (g/mL) potassium nitrate (CAS#7757-79-1), in distilled water. We adapted this PGM composition from Covey et al. 2010.

#### *In vitro* pollen growth assays

We collected pollen from individual flowers at anthesis on plants grown under the conditions described above, without heat-stress treatment, by gently vibrating open anther cones for 1 second with a modified electric toothbrush (Oral-B, DB4010) into a 1.5-mL microcentrifuge tube. Heat stress was first applied to the collected pollen during the subsequent in vitro germination and tube-growth assays. We added pre-warmed liquid PGM (control or heat-stress temperature) using a multi-channel pipette (BioHit, e1200), then resuspended the pollen by vigorous shaking in a custom tube rack. The pollen-PGM suspension was rapidly transferred to poly-D-lysine (CAS#26853-89-4)-coated glass-bottom 24-well plates (MatTek Corporation, P24G-1.5-13-F). Germinating pollen was maintained at 26°C (control) or 34°C (heat stress) using a heated stage insert (Biosciences Tools, TC-1-100i) supplemented with a custom-built stage heater consisting of polyimide heating film (12V 12W; 10mm x 93mm, Icstation) controlled by a W1209 temperature controller (HiLetgo). We selected 26 °C as the control temperature because it approximated the average ambient temperature of the microscope room. We selected 34 °C as the heat-stress temperature based on previous work demonstrating impaired pollen tube integrity in thermosensitive tomato cultivars at this temperature (Almasaud et al., 2026). Temperature was continuously monitored using a Raspberry Pi computer (Raspberry Pi Ltd) coupled to a Type T thermocouple (36 Gage PFA Wire, 40 inches; Evolution Sensors and Controls) (github.com/cedarwarman/pi_sensor).

We acquired images of germinating pollen every two minutes for two hours, beginning 15 minutes after PGM addition. We manually curated image series to include only fields containing 5-100 pollen grains, ensuring representative sampling while avoiding overcrowding. Each accession included in the analysis had at least seven usable replicates per condition. Replicate numbers varied among accessions because we excluded images with excessive pollen density, and plant or pollen sample loss occasionally limited the number of assays. The number of usable replicates per accession ranged from 7 to 25 at 26 °C (mean, 9.7) and from 7 to 19 at 34 °C (mean, 9.5).

#### Image pre-processing

Images were pre-processed as described in the workflow available at (github.com/cedarwarman/organize_and_process_microscope_images). We applied two procedures uniformly across the dataset. First, we normalized images to a consistent brightness level using the OpenCV Python library (v4.5.2). Second, we stabilized image sequences using the vidstab Python library (v1.7.4). We performed all image pre-processing on the University of Arizona High Performance Computing (HPC) cluster.

#### Image annotation

We used the online platform ‘Labelbox’ to generate annotated training data for computer vision models. We randomly selected images for manual annotation, ensuring that images from the same time-lapse sequence (i.e., images captured from one well during a single experimental run) were assigned exclusively to the training, validation, or test set. This prevented data leakage that could inflate model performance if images of the same pollen grains and tubes at different time points were used for both training and evaluation.

#### Image scoring

We annotated pollen grains with bounding boxes and classified them into four categories: germinated, ungerminated, burst, and aborted. A fifth category, ‘unknown_germinated’ was used when a visible pollen tube extended beyond the field of view, preventing us from determining whether it was intact or burst. Pollen grains in this category were excluded from all statistical analyses and plots.. We annotated pollen tube tips as points and classified them into three categories: tube_tip, tube_tip_burst, and tube_tip_bulging. In total, we labeled 1,000 images, yielding 79,253 annotated objects. Multiple annotators contributed to the labeling process and were randomly assigned images without knowledge of treatment conditions to minimize annotation biases.

#### Training and evaluation of convolutional neural networks

We evaluated publicly available object detection model architectures based on reported performance on the Collection of Common Objects (COCO) 2017 dataset. We selected CenterNet HourGlass104 for its high mean average precision (mAP) and relatively high processing speed (Newell et al. 2016; Zhou et al. 2019). We resized images to 1024 x 1024 pixels for training, evaluation, and inference. We trained and evaluated the model on the Jetstream2 system at Indiana University using an NVIDIA A100 GPU with 20 GB of memory. CenterNet HourGlass104 was implemented using the Google TensorFlow 2 Object Detection API (https://github.com/tensorflow/models/blob/master/research/object_detection/g3doc/tf2.md). We fine-tuned a model pre-trained on the COCO 2017 dataset on pollen image data using transfer learning. For training and evaluation, we randomly split the dataset into training (80%, n = 800 images) and evaluation (20%, n = 200 images) sets.

Detailed model configurations are available at (https://github.com/cedarwarman/pollen_cv/). Separate models were trained using all combinations of (i) images from camera 1, (ii) images from camera 2, (iii) combined camera images, (iv) pollen classes, (v) tube tip classes, and (vi) all classes combined (Supplemental Figure 2A). Models were trained for 30,000 steps (approximately 3 hours of wall time). We assessed performance using total loss, mAP, and average recall. We calculated a composite selection metric as a weighted sum of total loss and mAP. For each camera/class combination, we selected the training step that maximized this composite metric as the final model (Supplemental Figure 2B). We then performed final inference on the evaluation dataset for all models.

#### Object detection inference on the complete dataset

We performed inference on the complete dataset on the University of Arizona HPC using 4 NVIDIA V100 GPUs, each with 32 GB of memory. Approximately 300,000 images were analyzed in ∼19 hours of wall time. For each image, the model generated candidate bounding boxes for all classes along with associated confidence scores. We removed redundant detections using the TensorFlow implementation of the non-maximum suppression (NMS) algorithm. We determined the optimal confidence-score threshold by comparing model-predicted class counts from the validation set with ground-truth counts across confidence cutoffs from 0 to 1 in 0.01 increments (Supplemental Figure 2C). For each cutoff, we calculated the coefficient of determination (*R*²) between predicted and observed counts (https://github.com/cedarwarman/tomato_pollen_phenotyping/blob/main/R/compare_predictions_with_ground_truth.R). We selected the confidence threshold that maximized *R*² for downstream analyses.

#### Object tracking and class post-processing

We used the Bayesian Tracker (btrack) software package for multiple-object tracking and trajectory reconstruction (Ulicna et al. 2021). We implemented object tracking to determine which bounding boxes across consecutive frames in an image series correspond to the same object (https://github.com/cedarwarman/pollen_cv/blob/main/python/process_inference.py), enabling individual pollen grains and tube tips to be tracked throughout the time series. Btrack was customized to track pollen grains rather than mammalian cells, for which it was originally developed (https://github.com/cedarwarman/pollen_cv/blob/main/config/btrack/cell_config.json).

After object identification, we applied heuristic rules to refine pollen and tube tip class assignments over time. First, we interpolated objects across missing frames when detections were present in adjacent frames. Second, we corrected isolated misclassifications from the computer vision model when predicted state transitions were biologically implausible (e.g., a germinated pollen grain reverting to an ungerminated state, or a burst pollen grain reverting to an unburst state in a subsequent frame). Third, if a pollen grain was classified as aborted in >50% of frames within a track, we labeled the entire track as aborted; otherwise, we replaced aborted classifications with the preceding class assignment within the object’s time series.

After class correction, we linked individual tube-tip tracks to the nearest pollen grain track using the BallTree algorithm in scikit-learn (v1.0). Because time series were acquired using two cameras operating at different frame rates, sequences from camera 2 (66 frames per time series) were rescaled by linear interpolation to match the frame number of camera 1 (83 frames per time series) (https://github.com/cedarwarman/tomato_pollen_phenotyping/blob/main/R/plot_classes_over_time.R).

#### Comparison to hand counts

We evaluated model performance against ground-truth data by comparing the total number of bounding boxes for each class with counts obtained through manual annotation (https://github.com/cedarwarman/tomato_pollen_phenotyping/blob/main/R/compare_predictions_with_ground_truth.R). Models trained using data from both cameras were trained on 800 images. In contrast, models trained with data from a single camera (camera 1 or camera 2) were trained on 400 images. Test datasets consisted of 200 images for combined-camera models and 100 images for single-camera models. Test images were fully independent of training datasets, with no overlap in images or time series. For each class and camera combination, we compared manual counts and model predictions using linear regression models implemented in the R standard library and reported coefficients of determination (*R*²) (Supplemental Figure 2A).

#### Extracting phenotypes

For each accession, we calculated the percentage of objects in each class at each time point using class assignments predicted by the computer vision model after tracking-based correction. We computed burst integrals as the area under the bursting curve (burst percentage by time) using the trapezoidal method. We estimated pollen tube lengths from predicted track lengths.

## Statistical methods

Phenotypes from camera 1 and camera 2 images were evaluated for systematic biases by comparing accessions imaged with both cameras and by assessing potential effects of image acquisition date (https://github.com/cedarwarman/tomato_pollen_phenotyping/blob/main/R/run_statistics_on_cv_data.R). Experimental date, expressed as days since the start of the experiment, was included as a covariate. Because Heinz had more replicates than the other accessions, a subset of its replicates was randomly selected for statistical analysis. We corrected covariation between phenotypes and imaging data using Analysis of Covariance (ANCOVA) implemented in the R standard library (Supplemental Figure 3). We evaluated ANCOVA assumptions, including linearity, normality of residuals, and homogeneity of variance, for all phenotypes, including bursting rate, absolute bursting increase, burst integral, germination percentage, and pollen tube length. We calculated adjusted least-squares means using the emmeans R package (Lenth et al. 2024).

### Plotting and data organization scripts

We processed and organized data using the dplyr R package (Wickham et al. 2023), and generated figures using the ggplot2 R package (Wickham 2016). All code used to generate plots can be found at https://github.com/cedarwarman/tomato_pollen_phenotyping/.

## Results

### A diverse tomato panel for evaluating responses to elevated temperatures

We assembled a panel of 218 tomato accessions and their three close wild relatives: S. *lycopersicum* (171 accessions), S. *pimpinellifolium* (SP; 39 accessions), S. *cheesmaniae* (SC; 4 accessions), and S. *galapagense* (SG; 4 accessions). The panel was designed to capture the natural genetic diversity across cultivated varieties and the native range and regions associated with early domestication, including the Northern Andean region of South America, the Galápagos Islands, Central America, and Mexico (Razifard et al. 2020) (Figure 1A). We drew accessions from two previous panels encompassing broad phenotypic and genetic diversity (Mata-Nicolás et al. 2020, Alonge et al. 2020). We also included additional accessions previously reported to exhibit thermotolerance (Baggett 1985, Francesca 2022, Spormann 2023, Zhou 2017) (Supplementary Table 1).

**Figure 1:**
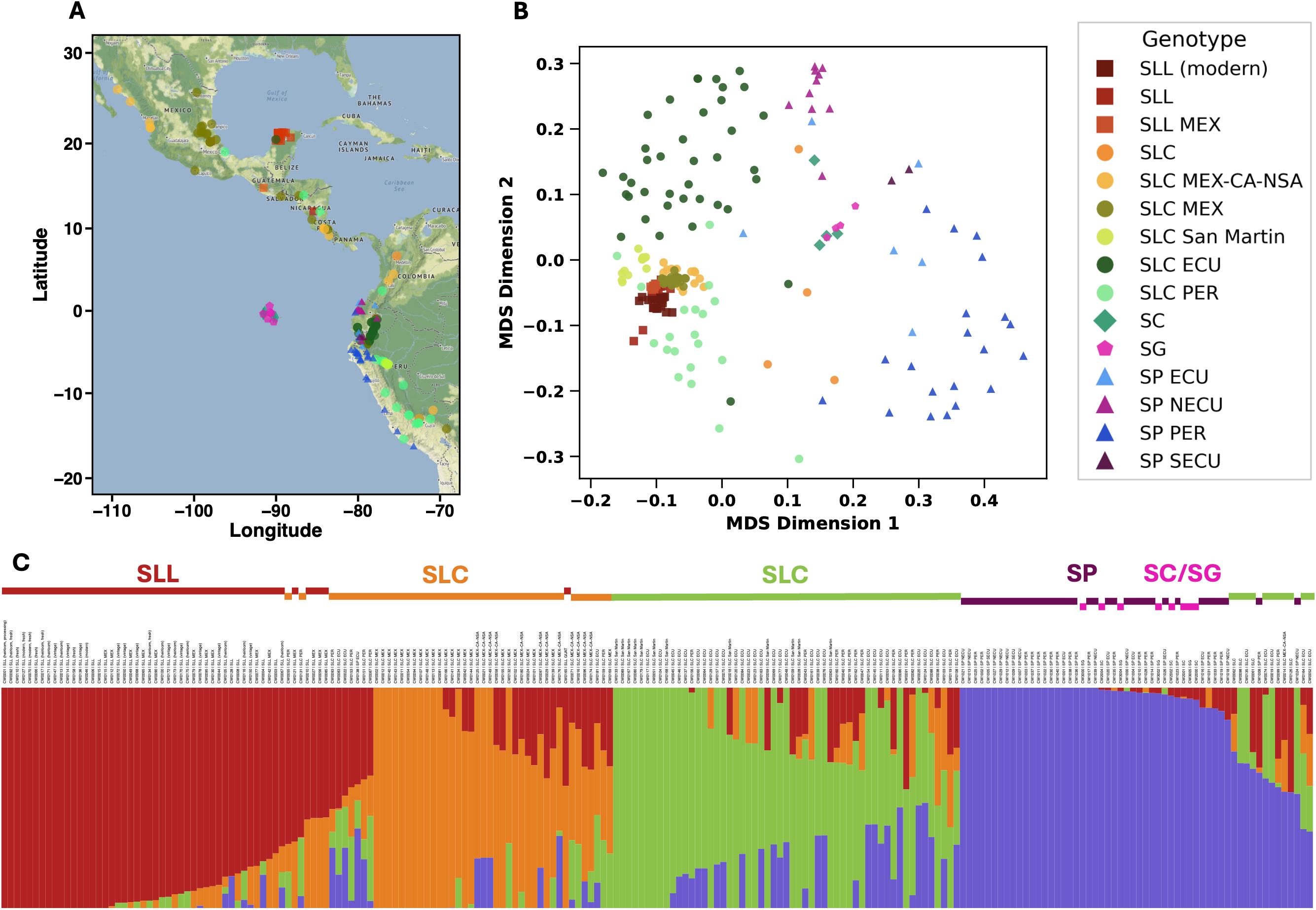
Diversity panel and population structure. A. Geographic distribution of collection locations of accessions included in the diversity panel. Sampling spans wild relatives from the native range in northwestern South America, early domesticated lineages from Central America and Mexico, and cultivated accessions representing modern diversity. The panel includes known thermotolerant varieties (Nagcarlang, Golden Nugget, and Tamulipas; Ouonkap et al., 2024). The shapes on the map indicate collection sites for five groups: *S. lycopersicum var. lycopersicum* (SLL; square), *lycopersicum var. cerasiforme* (SLC; circle), *S. pimpinellifolium* (SP; triangle), *S. cheesmaniae* (SC; diamond), and *S. galapagense* (SG; pentagon). The colors of the markers indicate the geographic distribution of accessions: MEX (Mexico), GUAT (Guatemala), MEX-CA-NSA (Mexico-Central America-Northern South America), ECU (Ecuador), NECU (Northern Ecuador), SECU (Southern Ecuador), PER (Peru). B. Multidimensional scaling (MDS) analysis of the diversity panel’s population structure. Points are colored by the same key as the geographic distribution. C. ADMIXTURE ancestry proportions (K = 4). Ancestry proportions inferred by ADMIXTURE at K = 4 with clusters largely aligning with the groups: (SP/SC/SG, purple; SLC ECU/SLC PER, green; SLL, red; SLC MEX, orange).

Of the 218 accessions, 208 had publicly available sequencing data that passed quality control and were retained for population structure analysis. Multidimensional scaling (MDS) of genome-wide pairwise identity-by-state (IBS) distances among these accessions, calculated from 3.1 million SNPs, revealed population structure broadly consistent with previous studies (Razifard *et al*., 2020). Cultivated *S. lycopersicum* var. *lycopersicum* (SLL) accessions formed a relatively compact cluster, whereas *S. lycopersicum* var. *cerasiforme* (SLC) accessions were more widely distributed (Figure 1B). The wild relatives SP, SC, and SG generally separated from the SLL accessions, with additional differentiation among SP accessions from different geographic regions, reflecting the isolation of their germplasm along the Andes and on Galápagos.

ADMIXTURE analysis at optimal *K*=4 further resolved ancestry patterns associated with species identity and geographic origin (Figure 1C). Wild SP, SC, and SG accessions formed a cluster distinct from the domesticated and semi-domesticated lineages geographically distributed primarily along the Pacific coast of South America and Galápagos (Pease *et al*., 2016; Razifard *et al*., 2020). Cultivated SLL accessions showed a strong genetic coherence and a geographic center in the Yucatán (Supplemental Figure 4). In contrast, the semi-domesticated, SLC accessions showed substantial admixture of two geographically structured groups of SLC-specific alleles and substantial admixed presence of alleles that predominate in either wild and/or cultivated accessions. One SLC ancestry cluster was concentrated in the Central Peruvian and Eastern Cordillera accessions, while the other SLC cluster was centered in Mexico and Central America. Several SLC accessions showed substantial representation across all four ancestry clusters, particularly at the Panamanian isthmus, Central Peru interchange zone, and far southern Peruvian Amazonia. Together, these analyses establish the genetic and geographic diversity of the panel, which we subsequently used to compare pollen germination, tube length, growth dynamics, and bursting at control and elevated temperatures.

### High-throughput microscopy and computer vision enable pollen phenotyping on a large scale

We designed a high-throughput microscopy system to image pollen grains and pollen tubes from accessions in the diversity panel under control and heat-stress conditions. Because accurate, scalable phenotyping underlies all downstream analyses, we describe the imaging, computer vision, and normalization pipeline in detail here. The system consisted of a Leica SP5 microscope with a programmable, temperature-controlled stage, integrated with computer-vision-based object detection and tracking pipelines for downstream analysis (Figure 2A; see Methods for details). We collected pollen from all accessions in the diversity panel and grew it *in vitro* in 24-well plates for 2 hours with continuous imaging. We precisely maintained temperatures at either control (26 °C) or heat-stress (34 °C) conditions. In total, we acquired approximately 300,000 pollen tube images. This scale of imaging enabled direct, quantitative comparison of pollen germination, tube growth, and bursting dynamics across genetically diverse accessions under heat stress.

**Figure 2:**
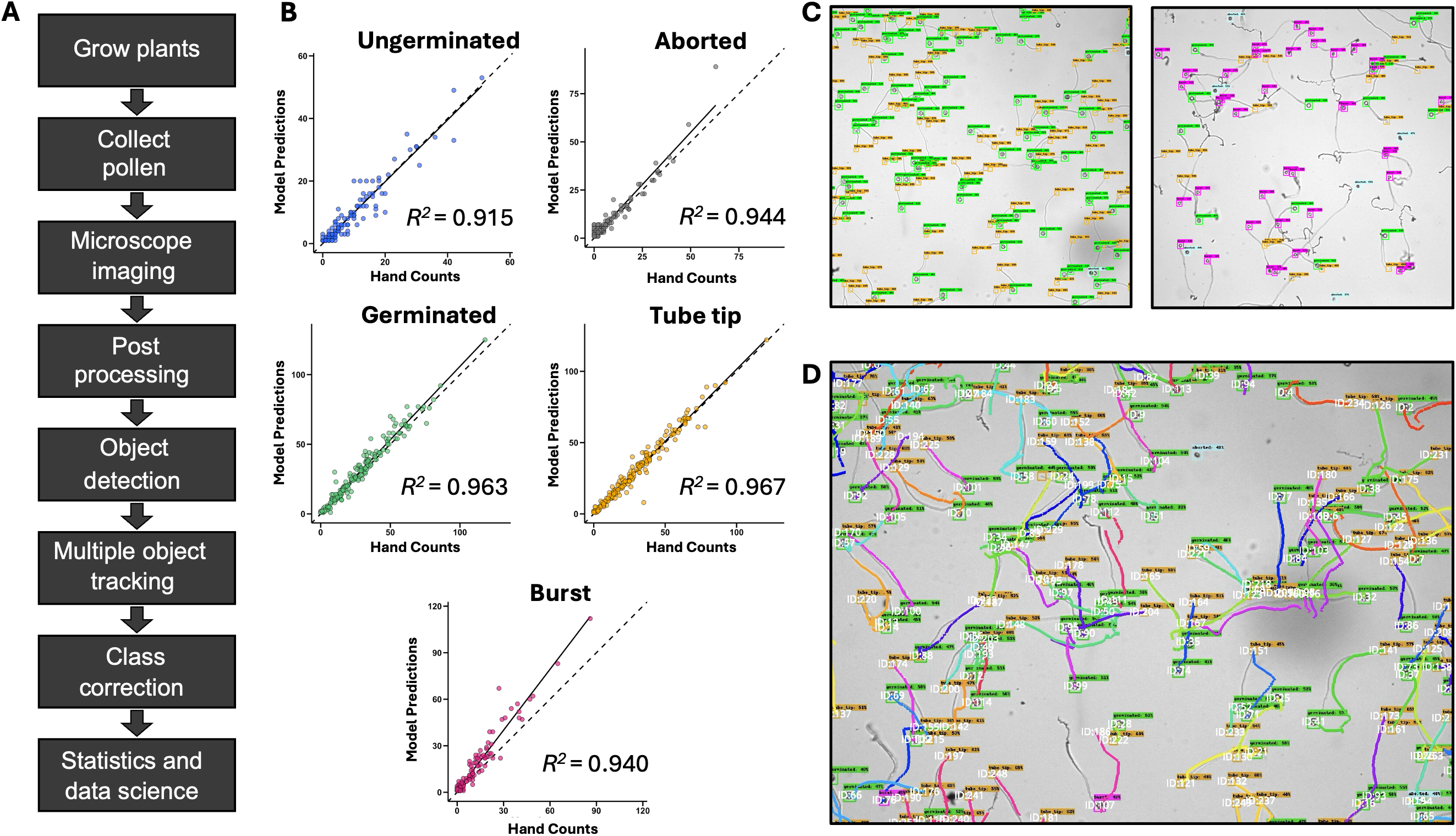
High-throughput phenotyping and model performance. A. Schematic overview of the high-throughput phenotyping pipeline used to quantify pollen traits. The workflow includes plant growth, pollen collection, bright-field microscopy imaging, image post-processing, object detection, multiple object tracking, class correction, and downstream statistical analysis. Detailed parameters and implementation are described in the Methods. B. Model performance benchmarking comparing automated predictions to manual counts across key pollen phenotypes. The model shows strong agreement for ungerminated, aborted, germinated, pollen tube tip, and burst classes, with high coefficients of determination (*R*² values shown), demonstrating the object detection model’s accuracy and robustness. C. Representative examples of object detection outputs. The model identifies and labels individual pollen phenotypes with bounding boxes: germinated pollen grains (green), pollen tube tips (orange), burst pollen grains (magenta), and aborted pollen grains (cyan). These examples illustrate the model’s ability to distinguish multiple phenotypic classes within complex images. The model accurately assigns labels even in crowded fields where pollen tubes cross paths, curve, or grow in different directions. D. Representative examples of multiple object tracking across time-lapse image sequences. Colored trajectories indicate the paths of individual pollen grains and growing tubes over time, enabling high-resolution quantification of pollen tube growth and behavior. Tracking remains robust even when pollen tubes intersect, overlap, or alter growth direction.

After image pre-processing, we manually annotated a subset of images to generate training data for computer vision model development. We reserved a separate, independently labeled image set for model evaluation. Four pollen classes were defined: (i) ungerminated pollen, lacking a visible tube or a tube shorter than the pollen grain radius; (ii) germinated pollen, with tubes longer than half the pollen grain diameter; (iii) burst pollen, characterized by tube rupture and release of cellular contents into the growth medium; and (iv) aborted pollen, which appeared smaller, darker, and lacked the mottled internal texture characteristic of healthy pollen. We also labeled tube tips at the distal ends of actively growing, unburst pollen tubes. These classes capture key cellular outcomes of successful or failed pollen tube growth that directly impact male fertility under thermal stress.

We trained an object-detection computer vision model (CenterNet HourGlass104) using the annotated training dataset and evaluated it using standard performance metrics (Supplemental Figure 1). After optimization, we performed inference on the held-out test dataset (Figure 2B). The model accurately detected all four pollen classes, including in images with high pollen density, debris, or burst pollen tubes. We assessed model performance by comparing predicted counts to manual counts from a randomly selected test image set. The correlation coefficient (*R²*) values were 0.915 for ungerminated pollen, 0.963 for germinated pollen, 0.940 for burst pollen, 0.944 for aborted pollen, and 0.967 for tube tips, indicating robust performance across all classes (Figure 2B). Figure 2C shows representative inference outputs. This accuracy enabled the detection of subtle but biologically meaningful differences in pollen performance among accessions.

After validation, we ran inference on the full image dataset, which required approximately 19 hours of computation and represented an estimated 2,700-fold speed increase over manual annotation. We aggregated counts of all detected objects for each accession at each time point. Because the object detection model did not link individual bounding boxes corresponding to the same pollen grain (for example, successive tube tip positions), we implemented multiple-object tracking to associate detections over time. We used a modified version of the Bayesian Tracker software (Ulicna et al. 2021) to generate time-resolved pollen tube trajectories (Figure 2D). We then used these tracks to refine pollen grain state classifications with biologically informed heuristics (see Methods), yielding an initial set of normalized pollen grain-class predictions and tube-length estimates. Linking detections over time enabled quantification of pollen tube growth dynamics at the individual-cell level rather than in static population snapshots.

Because we grew plants sequentially over several months in a greenhouse and sampled pollen many times, we examined phenotypes for potential covariation with experimental date (Supplemental Figure 2). We used burst percentage at 2 hours under heat stress (34 °C) as an initial diagnostic phenotype. Linear modeling revealed a significant negative correlation between burst percentage and experimental date, across all accessions (linear model *p <* 2.2 × 10^−16^) and within Heinz (control) replicates alone (linear model *p* = 5.6 × 10^−7^). These effects likely reflect physiological variation associated with plant growth conditions over time, underscoring the need to normalize data to reveal true genetic differences.

To account for the association between experimental date and pollen tube bursting this, we used ANCOVA models to adjust phenotypes for date-associated variation. For burst percentage at 2 hours and 34 °C, date was a significant covariate when accounting for accession identity (ANCOVA *p <* 2.2 × 10^−16^). A similar ANCOVA model for burst percentage at 2 hours under control conditions (26 °C) also identified collection date as a significant factor in the model (*p =* 3.6 × 10^−12^). Model assumptions were tested for all ANCOVA models (see Methods). Unbounded phenotypes, such as burst percentage increase or tube length, generally satisfy model assumptions more closely than bounded percentage-based traits. To quantify relative bursting dynamics while minimizing boundedness, we calculated a burst integral metric, defined as the area under the burst rate-versus-time curve (see Methods). We adjusted this and all other phenotypes using the least-squares means from the ANCOVA models, yielding the final phenotype values used in subsequent analyses presented here. This metric integrates the timing and extent of pollen tube failure, providing a biologically intuitive measure of thermotolerance.

### Heat stress negatively affects pollen tube germination and burst rates

We quantified pollen responses to temperature across all 218 accessions, allowing comparisons throughout the diversity panel. Heinz (SLL, Figure 1), a widely cultivated tomato accession reported to be relatively thermosensitive based on decreased pollen tube growth under heat stress (Ouonkap et al., 2024), and Tamaulipas (SLC MEX, Figure 1), reported to be a thermotolerant accession (Ouonkap et al., 2024), were examined in greater detail to illustrate contrasting growth dynamics. We plotted the mean percentage of each of the four pollen classes over time for all accessions, revealing several population-level patterns. First we assessed whether heat stress affected the proportion of aborted pollen grains. Pollen at the control temperature had a mean of 8.5% aborted grains, compared with 9.3% under heat stress; this difference was not significant (Supplementary Figure 5, paired *t*-test *p =* 0.071). We next examined how heat stress affected pollen germination. At control temperatures (Figure 3A), 50% of pollen grains germinated at a mean of 24.0 minutes after adding pollen growth medium, while under heat stress (Figure 3B), 50% germinated at a mean of 20.5 minutes. Thus, pollen grains germinated significantly faster under heat stress (Figure 3C, paired *t*-test *p <* 2.2 × 10^−16^).

**Figure 3:**
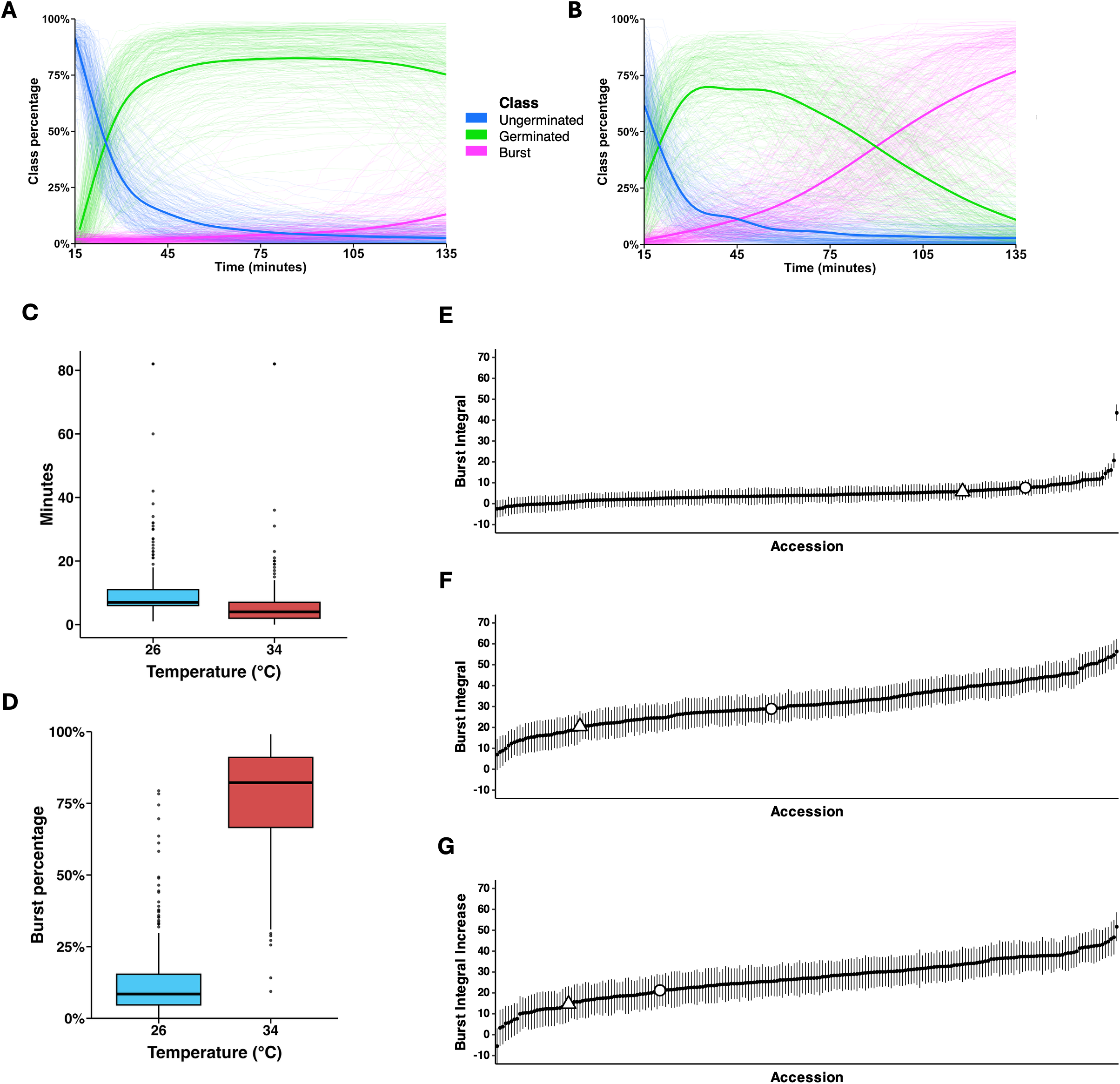
Heat stress increases pollen germination speed and tube bursting across the diversity panel. A-B. Temporal dynamics of pollen germination and bursting across all accessions at 26 °C (A) and 34 °C (B). Thin lines show accession-specific mean percentages over time, and thick lines show the mean across accessions at each time point. Colors indicate germinated pollen (green), ungerminated pollen (blue), and burst pollen tubes (magenta). Under control conditions, germination progresses with limited bursting. At elevated temperature, germination occurs earlier and bursting increases, with substantial variation among accessions. C. Comparison of time to 50% germination between control (26 °C) and heat stress (34 °C) conditions. Elevated temperature significantly reduces the time required to reach 50% germination, indicating faster germination under heat stress. D. Percentage of burst pollen tubes at 2 hours under control (26 °C) and heat stress (34 °C) conditions. Heat stress markedly increases bursting, reflecting compromised pollen tube integrity. E. Adjusted burst integral for each accession at 26 °C, summarizing cumulative bursting over time. Points represent individual accessions, and error bars indicate standard error. F. Adjusted burst integral for each accession at 34 °C. Values above the control indicate increased cumulative bursting under heat stress, with substantial variation across accessions. Error bars indicate standard error. G. Increase in adjusted burst integral for each accession (34 °C relative to 26 °C). This metric captures the extent of heat-induced bursting and reveals a wide distribution of responses across the diversity panel. Error bars indicate standard error. In E-G, the adjusted burst integral was determined by calculating the area under the curve of the percentage of burst pollen tubes over time, capturing the cumulative extent of bursting across the time course. In all three graphs, the x-axis represents individual accessions. The triangle indicates Tamaulipas and the open circle indicates Heinz.

In contrast to the modest acceleration in germination, heat stress profoundly affected pollen tube integrity. After 2 hours, pollen incubated at the control temperature showed a mean burst rate of 14.9%. In contrast, pollen exposed to heat stress showed a mean burst rate of 83.1%, a significant increase (Figure 3D, paired *t*-test *p <* 2.2 × 10^−16^).

To quantitatively capture bursting behavior over time, we calculated the area under the percent-burst curve using numerical integration. The “burst integral” reflects both the cumulative extent of pollen bursting and its temporal dynamics. Accessions that burst earlier and burst more rapidly yielded a higher burst integral than those that reached similar final burst percentages at later time points. Burst integrals varied widely among accessions. The mean burst integral was 4.63 at control temperature (Figure 3E), while increased significantly to 31.33 under heat stress (paired *t*-test, *p* < 2.2 × 10^−16^; Figure 3F). In addition to the increase in mean bursting, the variance in burst integrals was significantly higher at heat-stress temperatures (Levene’s test *p* < 2.2 × 10^−16^), indicating substantial variability in pollen tube response to heat stress.

In contrast, bursting responses were relatively uniform across accessions at the control temperature. To directly compare heat-induced bursting across accessions, we defined a relative metric termed “burst integral increase”, calculated as the burst integral at heat-stress temperature minus that at the control temperature. This metric summarizes each accession’s bursting response as a single, comparable phenotype (Figure 3G).

Most accessions in the diversity panel have not previously been evaluated for pollen thermotolerance. However, accessions with known temperature responses behaved as expected. Heinz exhibited a burst integral of 21.2, whereas Tamaulipas showed an apparently lower burst integral of 14.6. These values are consistent with their known heat-stress phenotypes and below the panel-wide mean increase in burst integral of 26.7 (Figure 3G).

### Heat stress affects pollen tube lengths across the tomato diversity panel

While the preceding analyses evaluated pollen phenotypes independently at each time point, measuring individual pollen tube growth required tracking the same tube over time. To address this challenge, we combined object detection and multiple object tracking to quantify individual pollen tube lengths throughout the time-lapse recordings for all accessions. Consistent with pollen germination and tube bursting phenotypes, pollen tube length varied widely across the diversity panel (Figures 4A, B). Overall, mean pollen tube length was significantly shorter under heat stress (773.5 µm) compared to control temperature (811.3 µm; paired *t*-test *p <* 2.2 × 10^−16^).

**Figure 4:**
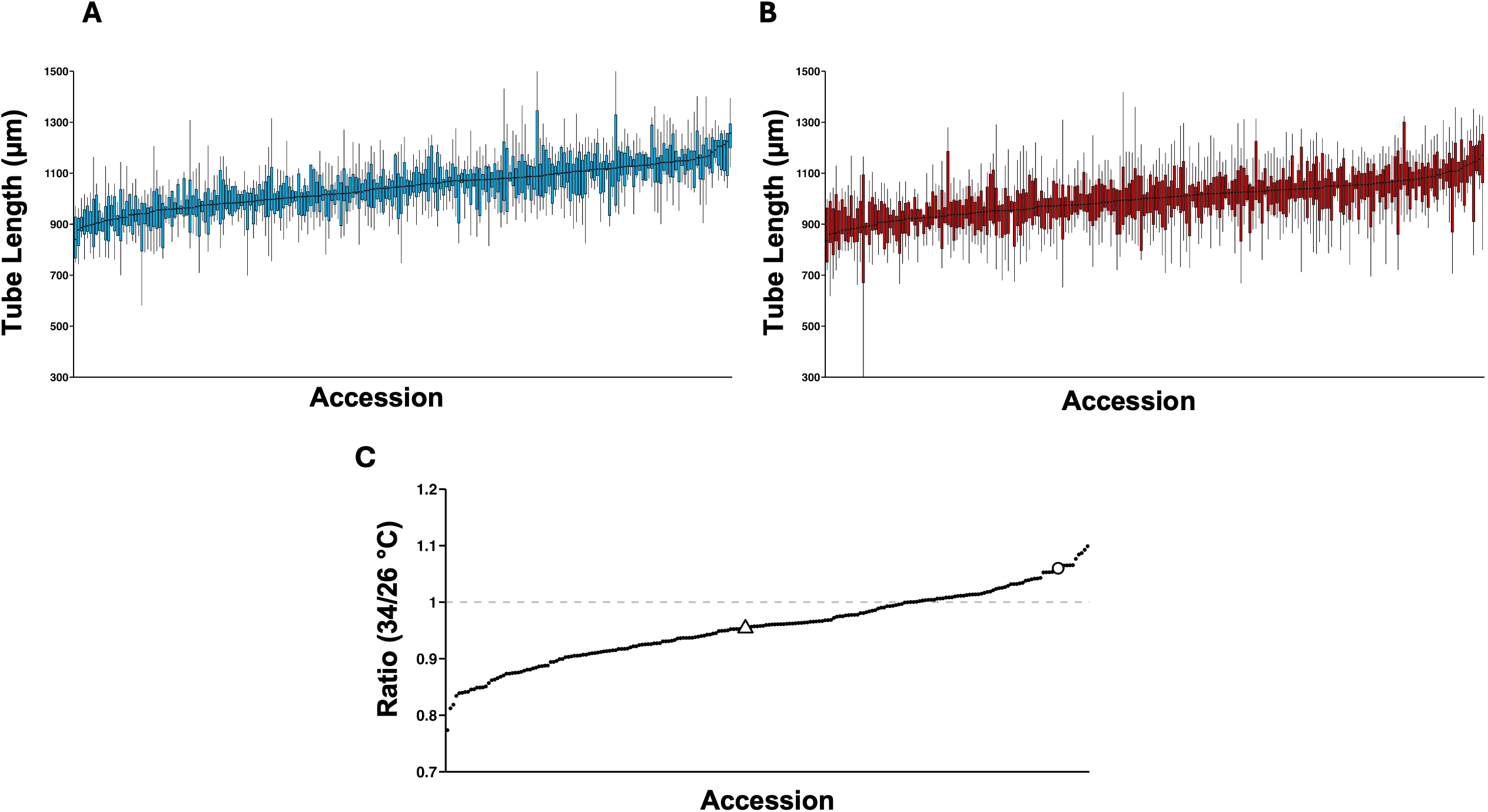
Accession-specific effects of heat stress on pollen tube length. A. Pollen tube lengths for each accession of the diversity panel at 26 °C, showing overall distribution and variation in growth under control conditions. B. Pollen tube lengths for each accession of the diversity panel at 34 °C, illustrating changes in growth under heat stress across the diversity panel. C. Ratio of mean pollen tube length at 34 °C to that at 26 °C for each accession. Tamaulipas (circle) and Heinz (triangle) are highlighted as previously characterized thermotolerant and relatively thermosensitive reference accessions, respectively (Ouonkap et al., 2024). The dotted horizontal line indicates a ratio of 1, representing equal mean lengths at both temperatures. In (A) and (B), boxes span the first to third quartiles (interquartile range, IQR), and the line within each box indicates the median. Whiskers extend to the smallest and largest observations within 1.5 × IQR below the first quartile and above the third quartile, respectively.

Despite this overall reduction in tube length, responses varied substantially across accessions. To quantify accession-specific responses, we calculated the ratio of mean pollen tube length at 34 °C to that at 26 °C (Figure 4C). Ratios below 1 indicate reduced pollen tube elongation under heat stress, whereas ratios above 1 indicate equal or enhanced tube elongation at elevated temperature. Most accessions (72%) exhibited shorter tubes at 34 °C, while a smaller subset (28%) maintained or increased tube length. Representative accessions reflected these contrasting phenotypes. Heinz, a moderately thermosensitive accession, exhibited reduced pollen tube lengths under heat stress (ratio = 0.90), whereas Tamaulipas, a thermotolerant accession, produced longer pollen tubes at elevated temperature (ratio > 1). Together, these results suggest substantial natural variation in pollen tube growth response to elevated temperature across the tomato diversity panel.

### Heat stress reshapes pollen tube growth dynamics and reveals accession-specific growth speed distributions

In addition to final tube length (Figure 4A-C), we examined pollen tube growth dynamics by quantifying growth speed over time for all accessions at control and heat-stress temperatures. In general, at 26 °C, mean pollen tube growth speed initially declined slightly, then increased to a peak at approximately 90 minutes before declining modestly toward the end of the assay (Figure 5A). Growth trajectories varied substantially among accessions at 26 °C, with many showing increasing growth speeds after an initial decline during the first 20–30 minutes (Figure 5B). In general, at 34 °C, mean growth speed was initially higher than at 26 °C, increased to a peak at approximately 40–50 minutes, and then declined through the remainder of the assay (Figure 5A). Individual accessions appeared to differ in the timing and magnitude of these changes. As with tube length, growth-speed responses to heat stress varied across the diversity panel.

**Figure 5:**
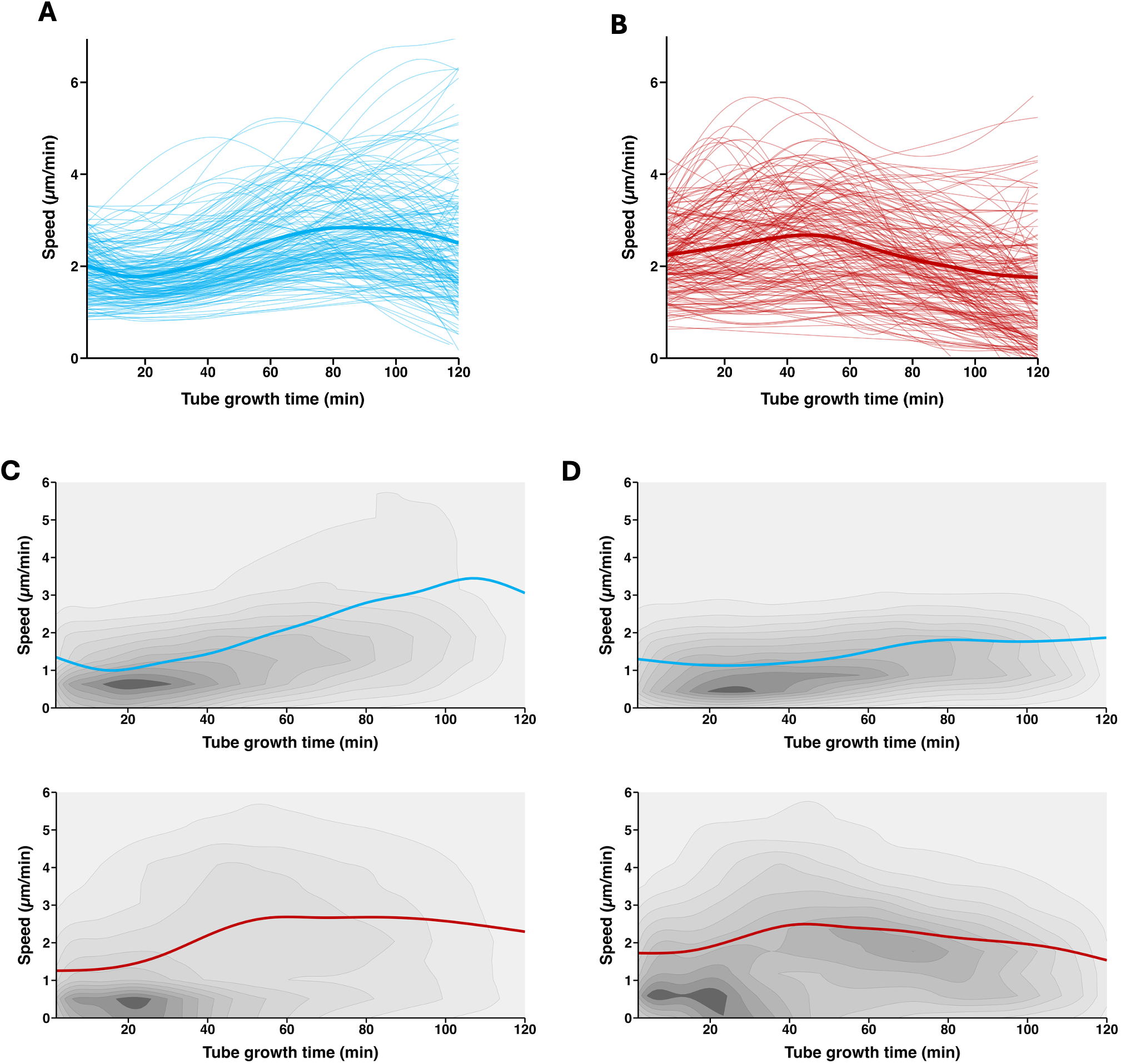
Accession-specific effects of heat stress on pollen tube growth speed. A. Pollen tube growth speed over time for all accessions at 26 °C. Individual lines represent accession-specific trajectories, with an overall trend toward increasing growth rate during the early and mid phases of growth. B. Pollen tube growth speed over time for all accessions at 34 °C. Growth is initially faster under heat stress but declines over time in many accessions, reflecting altered growth dynamics. C. Growth speed distribution for the thermosensitive accession Heinz at 26 °C (top) and 34 °C (bottom), showing a narrower distribution skewed toward lower growth speeds. D. Growth speed distribution for the thermotolerant accession Tamaulipas at 26 °C (top) and 34 °C (bottom), showing a broader distribution with a sustained subpopulation of faster-growing pollen tubes under heat stress (34 °C).

To examine accession-specific growth dynamics, we compared pollen tube growth rates in Heinz and Tamaulipas accessions. Under control conditions (26 °C), Heinz pollen tubes showed a sustained increase in growth speed through most of the assay, whereas Tamaulipas exhibited a more modest increase followed by relatively stable growth. These trajectories broadly matched those reported at 28 °C using TubeTracker (Ouonkap et al., 2025), despite the temperature difference; at 34 °C, Heinz growth speed increased to a plateau around 60 minutes before gradually declining. Tamaulipas reached its maximum growth speed earlier, around 40 minutes, then gradually declined over the remainder of the assay. Thus, elevated temperature altered the growth trajectories of both accessions, which reached peak growth speed at different times before gradually slowing.

We next extended this analysis to compare growth-speed distributions over time under control and heat-stress conditions, revealing contrasting responses in Heinz and Tamaulipas. Under control conditions, both Heinz and Tamaulipas showed a single predominant concentration of pollen tube growth speeds, with no clearly separated subpopulations (Figures 5C and 5D). At 34 °C, Heinz was dominated by slower-growing tubes, with relatively few fast-growing tubes persisting over time (Figure 5C). By contrast, Tamaulipas showed a broader distribution, with an apparent separation between slower-growing tubes and a subset of faster-growing tubes that persisted throughout much of the 2-hour assay (Figure 5D). These contrasting distributions reveal apparent differences in pollen responses to heat that endpoint tube length alone does not fully describe. Whether these apparent subpopulations are statistically distinct, and whether similar patterns occur in other accessions, will require further analysis.

## Discussion

### A scalable framework for population-level phenotyping of pollen tube growth

Automated image analysis has expanded the scope of quantitative analysis of pollen traits, enabling staining-based viability assessment (Ascari et al., 2020; Tan et al., 2022), measurements of germination and tube length (Zhang et al., 2023; Yamazaki et al., 2023), and time-lapse analysis of germination timing, elongation rate, and tube integrity (Ouonkap et al., 2025). Automated pollen viability screening has also supported investigations of the genetic basis of reproductive thermotolerance in maize (Li et al., 2026). Our phenotyping platform builds on these advances by integrating automated microscopy, object detection, multiple-object tracking, and temporal analysis within a single workflow. Applied to over 300,000 images from 218 tomato and wild-relative accessions under control and heat-stress conditions, this platform transforms a traditionally low-throughput pollen tube growth assay into a phenomics-scale approach for quantifying multiple pollen performance traits across a large and diverse germplasm sample.

A key advantage of this workflow is its integration of high-throughput image acquisition with complementary analytical tasks. Object detection scores pollen phenotypes independently in each image, while multiple-object tracking links individual pollen tubes across successive frames to quantify tube elongation over time. Together, these approaches characterize pollen performance at both accession and individual-tube scales while reducing the need for manual scoring. Agreement between automated predictions and manual annotations supports the accuracy of the measured traits. This validation is particularly important across genetically diverse material, where differences in pollen morphology and growth behavior may challenge consistent detection and measurement. These capabilities provide a validated platform for population-scale comparisons of pollen performance under control and heat-stress conditions.

### Heat stress accelerates germination but compromises pollen tube integrity

The consistently low proportion of aborted pollen suggests that our plant growth conditions generally supported normal pollen development. We applied heat stress only to mature pollen during *in vitro* germination and tube growth. Future applications could examine pollen collected after developmental heat treatments, for example, following the approach of Begcy et al., (2019), who applied transient heat stress at the tetrad stage in maize and observed impaired mature pollen function. Under such conditions, quantifying aborted classes could inform assessments of heat stress effects on development alongside germination and tube growth.

Our results reveal that heat stress affects distinct stages of pollen tube growth differently. Accelerated germination was accompanied by increased pollen tube bursting in most accessions. These findings are consistent with Almasaud et al. (2026), who observed accelerated germination at elevated temperature across six tomato cultivars, together with increased rupture in thermosensitive Heinz and VF36. In that study, maintenance of pollen tube integrity distinguished thermotolerant from thermosensitive cultivars, rather than elongation rate alone. Our population-scale analysis extends these observations across a broad panel of cultivated tomato and wild relatives, and further demonstrates germination and subsequent tube integrity are distinct components of pollen thermotolerance.

Heat stress accelerated germination but significantly increased pollen tube bursting in most accessions, suggesting disruption of processes that maintain cellular integrity during growth. Premature rupture prevents sperm delivery, making tube integrity an important component of reproductive thermotolerance. Almasaud et al. (2026) linked heat-induced integrity loss to altered RALF signaling and pectin distribution. However, loss of SlRALF1 in thermosensitive Heinz improved tube integrity and increased seed production in crosses using heat-exposed pollen. These findings provide a partial mechanistic context for the bursting variation in our panel.

### Natural variation in pollen thermotolerance is extensive across tomato germplasm

Population-scale phenotyping revealed apparently extensive natural variation in pollen thermotolerance across the tomato diversity panel. Accessions differed in heat-induced germination rates, pollen tube length, growth dynamics, and bursting. Notably, accessions previously characterized as thermotolerant or thermosensitive based on fruit and seed set under heat stress (Ouonkap et al., 2024) showed i*n vitro* pollen performance at high temperature consistent with their *in vivo* classification. This concordance provides independent support for the phenotyping approach and the biological relevance of the measured pollen tube growth traits.

The variation in pollen performance under heat stress among cultivated tomato and wild-relative accessions highlights the substantial genetic diversity available for improving reproductive resilience to heat stress. Differences in pollen performance may reflect genetic variation shaped by adaptation to natural environments or domestication and breeding. The quantitative phenotypes generated in this study provide a valuable resource for determining genetic associations between genetic variation and pollen thermotolerance. More broadly, pollen performance traits are rapidly screenable intermediate phenotypes linking genetic variation to reproductive resilience under heat stress, and, ultimately, crop productivity in warming environments. Additional information on heat impacts on reproductive performance across the accession panel will require evaluation of fruit and seed set under heat stress.

### Dynamic phenotypes reveal aspects of thermotolerance missed by endpoint measurements

Endpoint measurements such as final germination percentage, tube length, or bursting rate summarize pollen performance but do not resolve how these outcomes develop over time. By continuously monitoring pollen behavior throughout the assay, our framework captures temporal patterns that endpoint measurements would miss. Heat stress altered the extent and timing of pollen germination and tube bursting. These temporal offsets are not measurable when data collection is restricted to a single time point.

The burst integral metric introduced here innovates an integration of the extent and temporal progression of pollen tube bursting over the observation period (2 hours in this study). For accessions with similar final burst percentages, earlier bursting produces higher integral values than delayed bursting, distinguishing responses that an endpoint comparison would treat as equivalent (Supplemental Figure 6). Because premature tube rupture is likely more detrimental to fertilization success than rupture occurring later in pollen tube growth, this metric may provide a more biologically meaningful representation of pollen fitness than endpoint measurements alone. Dynamic phenotypes such as burst integral therefore offer a promising framework for quantifying thermotolerance and may prove especially powerful for identifying underlying genetic determinants of thermotolerance.

### Thermotolerant accessions maintain distinct pollen growth dynamics under heat stress

Beyond resolving the timing of pollen tube bursting, tracking individual tubes revealed how heat stress altered growth dynamics within and among accessions. Analysis of individual pollen tube growth trajectories revealed differences in heat responses that considering tube length alone at the end of the 120-minute imaging period would have missed. Although most accessions exhibited reduced tube length at elevated temperature, their growth rates followed different trajectories over time. Thermosensitive accessions showed a progressive decline in rapidly growing pollen tubes and a shift toward slower growth-speed distributions. In contrast, thermotolerant accessions maintained broader distributions, retaining subpopulations of pollen tubes that sustained elongation throughout the assay. These patterns suggest that the persistence of an actively growing pollen subpopulation in an accession may be an important component of pollen thermotolerance.

This finding is consistent with reports of pollen subpopulations with contrasting ROS levels, in which lower ROS levels were associated with greater thermotolerance (Rutley *et al*., 2021; James *et al*., 2026). The parallel raises the possibility that differences in ROS homeostasis contribute to the growth heterogeneity observed in our assay.. Maintaining cell wall integrity, membrane stability, and efficient stress-response pathways could also support continued elongation under heat stress. Together, these findings show that growth-rate distributions reveal variation both among accessions and among individual pollen tubes, complementing accession-level summaries of pollen performance.

### Future applications and opportunities for gene discovery

The phenotyping framework presented in this study provides a foundation for identifying genetic determinants of pollen thermotolerance. Combining the quantitative traits measured using this phenotyping system across this diversity panel with available genomic resources will enable genome-wide association studies to identify loci and variants associated with germination, tube elongation, and integrity maintenance under heat stress. Candidate genes and genomic loci identified using these methods can subsequently be tested through gene editing, transgenic approaches, and targeted physiological assays.

By examining pollen directly under controlled conditions, *in vitro* assays help distinguish pollen responses from temperature effects on surrounding floral tissues and whole-plant physiology. Pollen’s haploid state also offers opportunities to investigate allelic effects expressed in the male gametophyte (Johnson et al., 2004), although distinguishing these from parental sporophyte effects will require additional experiments. By identifying pollen-specific alleles involved in thermotolerance, this system also allows researchers to examine this component of reproductive resilience separately from the broader complexity of whole-plant heat tolerance.

This phenotyping system could be extended to pollen from other flowering plants and environmental stress applications, provided that culture conditions, imaging procedures, and analytical models are validated for each application. Potential extensions include pollen responses to drought, salinity, UV stress, or chemical perturbations. Because pollen tube growth is a sensitive single-cell component of reproductive fitness, large-scale phenotyping of pollen populations offers a powerful way to connect cellular phenotypes with agronomically important traits. This system provides a relatively inexpensive first step for screening pollen thermotolerance under controlled conditions, reducing the need to evaluate every accession in resource-intensive field trials. Accessions with promising pollen performance can then be prioritized for testing pollen tube growth in the pistil, fruit and seed set, and yield under heat stress. Integrating high-throughput pollen phenotyping with modern genomics and breeding approaches may accelerate the development of crops that maintain reproductive success under increasingly challenging climatic conditions.

## Conclusions

This study establishes an integrated framework combining automated microscopy, computer vision, and individual pollen tube tracking to quantify pollen function across genetically diverse populations over time. Application to 218 tomato and wild-relative accessions revealed that elevated temperature accelerated pollen germination, compromised tube integrity through increased bursting, and altered elongation, with substantial variation among accessions. Time-resolved measurements distinguished the progression of germination,bursting, and growth-speed distributions that endpoint measurements could not fully describe. These findings demonstrate the value of evaluating multiple, dynamic traits when characterizing pollen responses to heat. The framework and resulting phenotypic dataset provide a foundation for identifying genetic determinants of pollen thermotolerance and selecting contrasting accessions for mechanistic studies. As an efficient initial screening approach, the system could also rapidly prioritize germplasm for subsequent evaluation of pollen performance in the pistil, fruit and seed set, and yield under heat stress. Together, these advances make dynamic pollen behavior accessible to population-scale investigation of reproductive responses to environmental stress.

## Data Availability Statement

All data supporting this study’s findings are publicly available. Processed datasets and code used for data analysis are available in a GitHub repository. Where applicable, specific links to datasets, code, and analysis pipelines are provided in the Methods section. Additional data and resources are provided within the article and its Supplementary Information.

## AI Code Disclosure

AI-assisted tools were used in a limited, supportive capacity to improve efficiency, specifically to check and polish code used in data analysis and to help finalize figures. No AI tools were used to generate original scientific content, analyze data, or produce results or conclusions. The authors developed, verified, and validated all analyses, interpretations, and code used in this study.

## Author Contributions

CW and RP conceptualized the study; CW and JP performed population genomic analyses; CW, SM, and CJP performed the experiments; BG generated the map showing the locations where the community collected the accessions used in the diversity panel; KT developed some code for data analysis; CW and CJP performed data curation and formal analysis with input from SES, who provided statistical consultation and validation; RP supervised the project and managed execution; CW, CJP, JP, SES, and RP wrote the manuscript with input from other authors.

## Acknowledgements

Thank you to Yaire Gutierrez, Jorge Munoz, Xander Nelson, Christopher Nacion, and Katrina Zearley for technical assistance. Thank you to University of Arizona High Performance Computing and Indiana University Jetstream2 for computational resources. Thank you to Dr. Fernando Rios and his team at ReDATA, University of Arizona Libraries Data Cooperative, for publicly hosting the image datasets reported in this study. Thank you to the C.M. Rick Tomato Genetics Resource Center, the USDA National Plant Germplasm System, and the labs of Dr. Esther van der Knaap and Dr. Zachary Lippman for tomato germplasm. We acknowledge the greenhouse support of Arturo Baez and his team at the University of Arizona Campus Agriculture Center. We thank Drs. Mark Johnson, Gloria Muday, Ann Loraine, Robert Reid, and James Pease, along with their laboratory members, for their valuable suggestions and discussions throughout this study during NSF IOS-1939255 project meetings. This work was funded by National Science Foundation (NSF) grants-NSF IOS-1939255 (RP and JP), NSF IOS-2109832 (CW), the United States-Israel Binational Agricultural Research and Development Fund (BARD-IS-5510-22) (RP and CJP), the Biochemistry and Molecular and Cellular Biology (BMCB) Predoctoral Training Grant (NIH-T32GM136536) (CJP) and NSF Center for Research on Programmable Plant Systems (DBI-2019674) (CJP and RP).

## Supplemental Figure Legends

**Supplemental Figure 1. Seedling, floral, and fruit diversity across the tomato accessions used in this study.** A. Montage of seedling images illustrating variation in plant stature, leaf morphology, and growth habit. B. Montage of flowering shoots illustrating variation in floral and inflorescence morphology. C. Montage of fruit images illustrating variation in size, shape, color, and arrangement on the truss. Black-bordered insets in each panel show enlarged examples from the corresponding montages to illustrate details. A color reference chart is visible in the seedling inset (A), and a ruler provides a size reference in the fruit inset (C).

**Supplemental Figure 2. Model training evaluation, camera comparison, and confidence threshold selection.** A. Comparison of model performance using individual camera datasets versus combined camera training. The mean coefficient of determination between the training and validation sets is shown for each pollen phenotype class (ungerminated, germinated, burst, aborted, and tube tip), demonstrating the impact of the training data source on model accuracy. B. Model training metrics across training iterations using combined camera datasets. We evaluated models trained on all classes, pollen-specific classes, and tube tip classes. Metrics shown include precision, mean average precision (mAP), average recall (AR), and a combined performance score. We used these curves to assess model convergence and identify the optimal training checkpoint. C. Model performance across a range of confidence thresholds for each phenotype class. We used these curves to determine the optimal model confidence threshold that balances accuracy across classes.

**Supplemental Figure 3. Correction of temporal effects on pollen tube bursting measurements.** A. Effect of experimental date on pollen tube bursting at 34 °C. Each point represents the percentage of burst pollen tubes at 60 minutes for an individual assay, plotted over time. Data is colored by the imaging camera. A negative trend over time indicates a significant date effect on bursting measurements (R² = 0.1739, *p <* 2.2 × 10 ¹), whereas camera effects alone are not a major source of variation. B. Adjustment of burst measurements using ANCOVA. Raw and adjusted burst integral values at 34 °C are shown across experimental days. Adjusted values represent least-squares means corrected for date effects, reducing temporal bias and enabling more accurate comparison across accessions and experiments.

**Supplemental Figure 4. Geographic distribution and genetic structure of ADMIXTURE ancestry clusters in the tomato diversity panel.** A. Collection sites of georeferences accessions, each shown as a pie chart of its ADMIXTURE ancestry proportions at K = 4. Cluster colors: 1 (red), predominantly SLL; 2 (orange), SLC from Mexico and Central America; 3 (green), SLC from Ecuador and Peru; 4 (purple), SP, SC, and SG. Each cluster is centered in a distinct geographic region, and admixed accessions occur predominantly outside these centers. Accessions sharing identical coordinates are arranged in a ring around the shared site. B. Metric multidimensional scaling of pairwise identity-by-state genetic distances among 208 accessions, with each accession shown as a pie chart of its ADMIXTURE ancestry proportions at K = 4. Colors in (A). In both panels, ancestry proportions below 1% are omitted for clarity.

**Supplemental Figure 5. Transient heat stress does not affect the proportion of aborted pollen grains.** (A) The proportion of aborted pollen grains increases slightly over the assay time course, independent of temperature. Mean proportion of aborted pollen grains from 15 to 135 minutes after adding pollen growth medium, at the control temperature (26 °C, blue) and under heat stress (34 °C, orange). Lines show the mean across accessions, and shaded ribbons show 95% confidence intervals. Comparing accession means from the first five frames (15–21 min) with the last five (129–135 min), the aborted proportion rose by a mean of 0.9 percentage points at both 26 °C (8.0% to 9.0%; n = 226 accessions; paired t-test, *p* = 8.3 × 10 ^12^) and 34 °C (8.6% to 9.5%; n = 226 accessions; *p* = 1.1 × 10 ^8^). The magnitude of this increase did not differ between temperatures (n = 226 accessions with both; paired t-test, *p* = 0.95). (B) The proportion of aborted pollen grains in 226 accessions of the diversity panel, assayed at the control temperature (26 °C) and under heat stress (34 °C). Each point is the mean proportion of aborted grains for one accession, with each well summarized as the median across all time points of the 2-hour assay. Violins show the distribution of accession means; boxplots show the median and interquartile range, with whiskers extending to 1.5 times the interquartile range. Aborted grain proportions did not differ between temperatures (paired t-test, *p* = 0.081; n.s., not significant).

## Supplemental Tables

Supplemental Table S1. Accession information for the tomato diversity panel used in this study.

Supplemental Table S2. Genomic data quality control and variant calling.

