## Supplementary figures and images for "High-throughput phenotyping reveals natural variation in tomato pollen responses to heat stress"

### Supplemental Figure S1

#### Slide 1
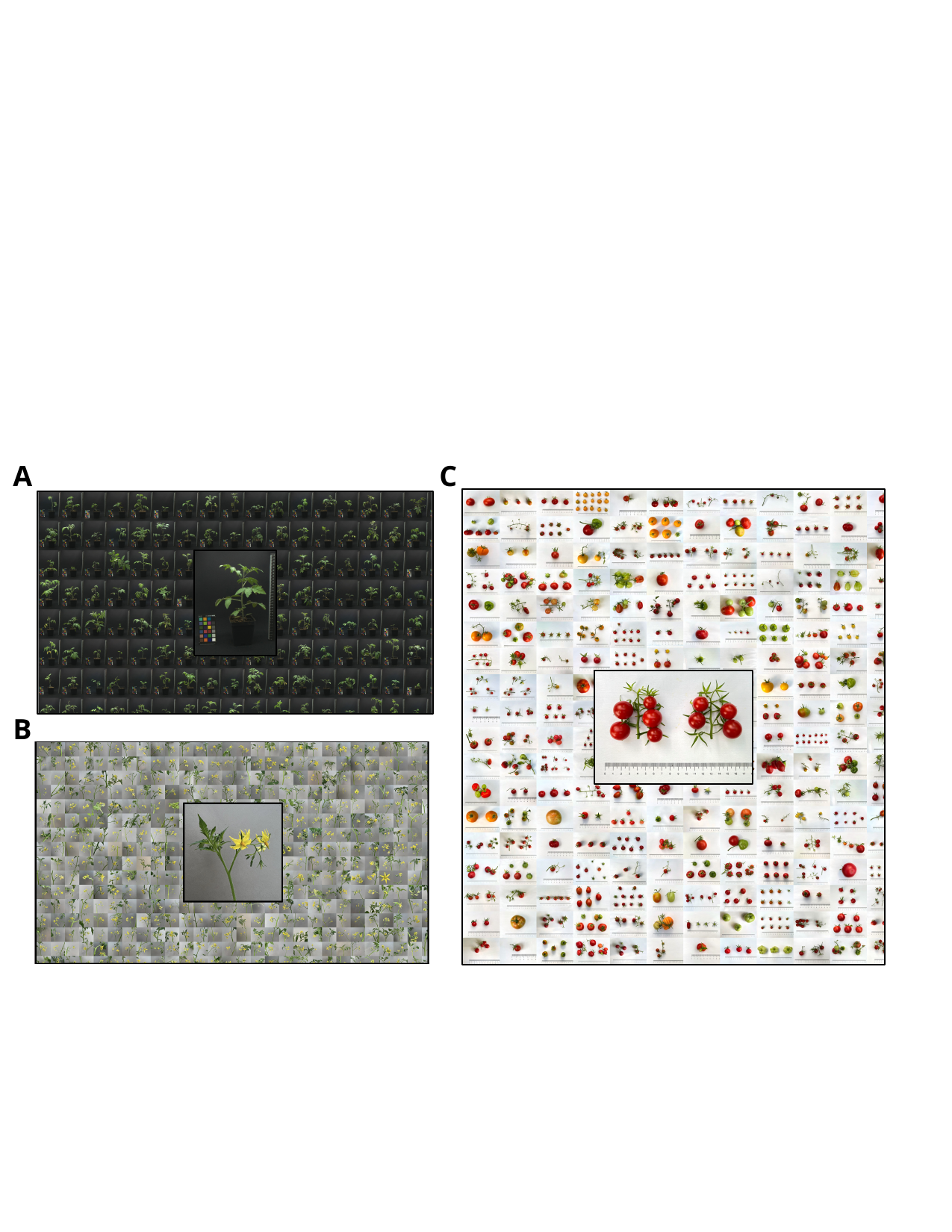

A
C
B

### Supplemental Figure S2

#### Slide 1
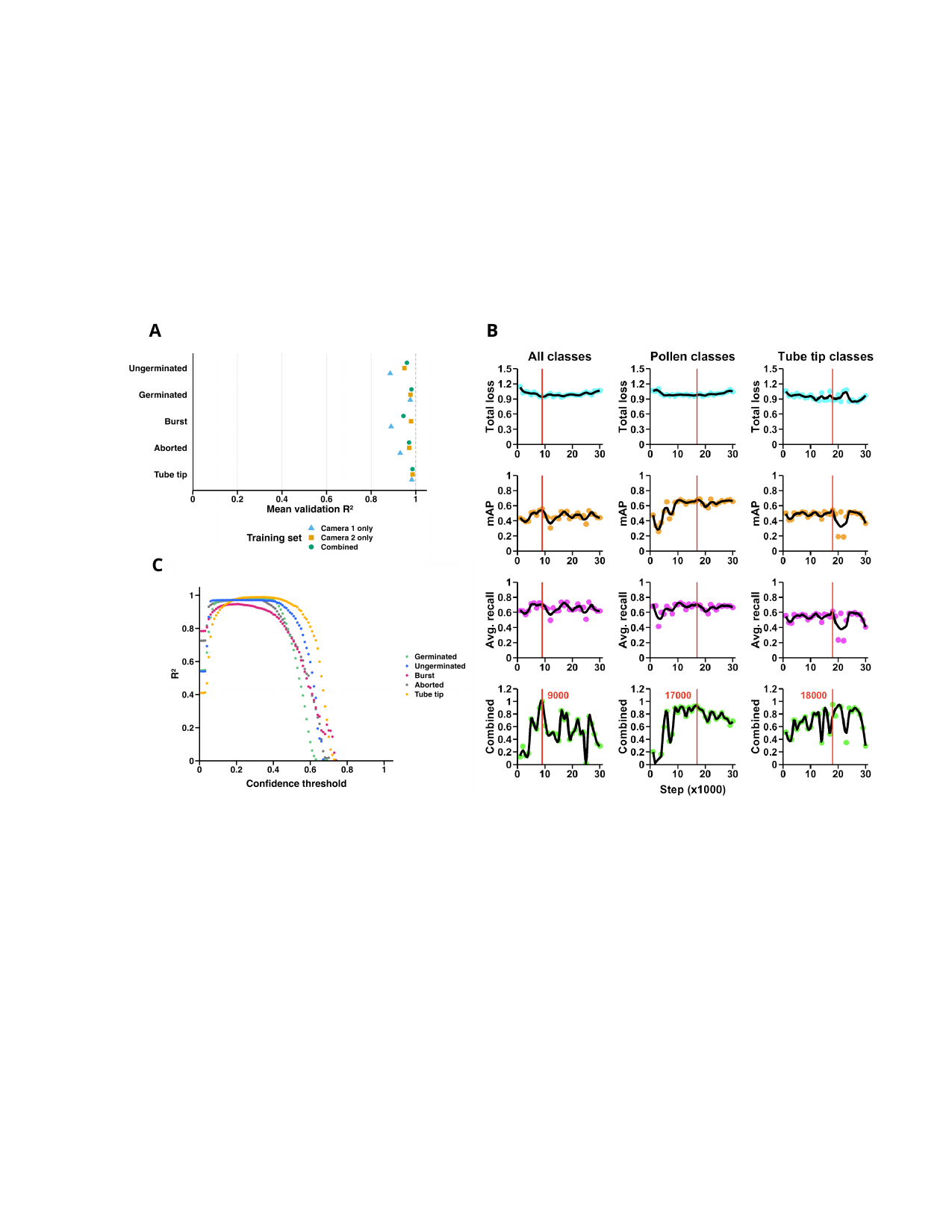

A
B
C

### Supplemental Figure S3

#### Slide 1
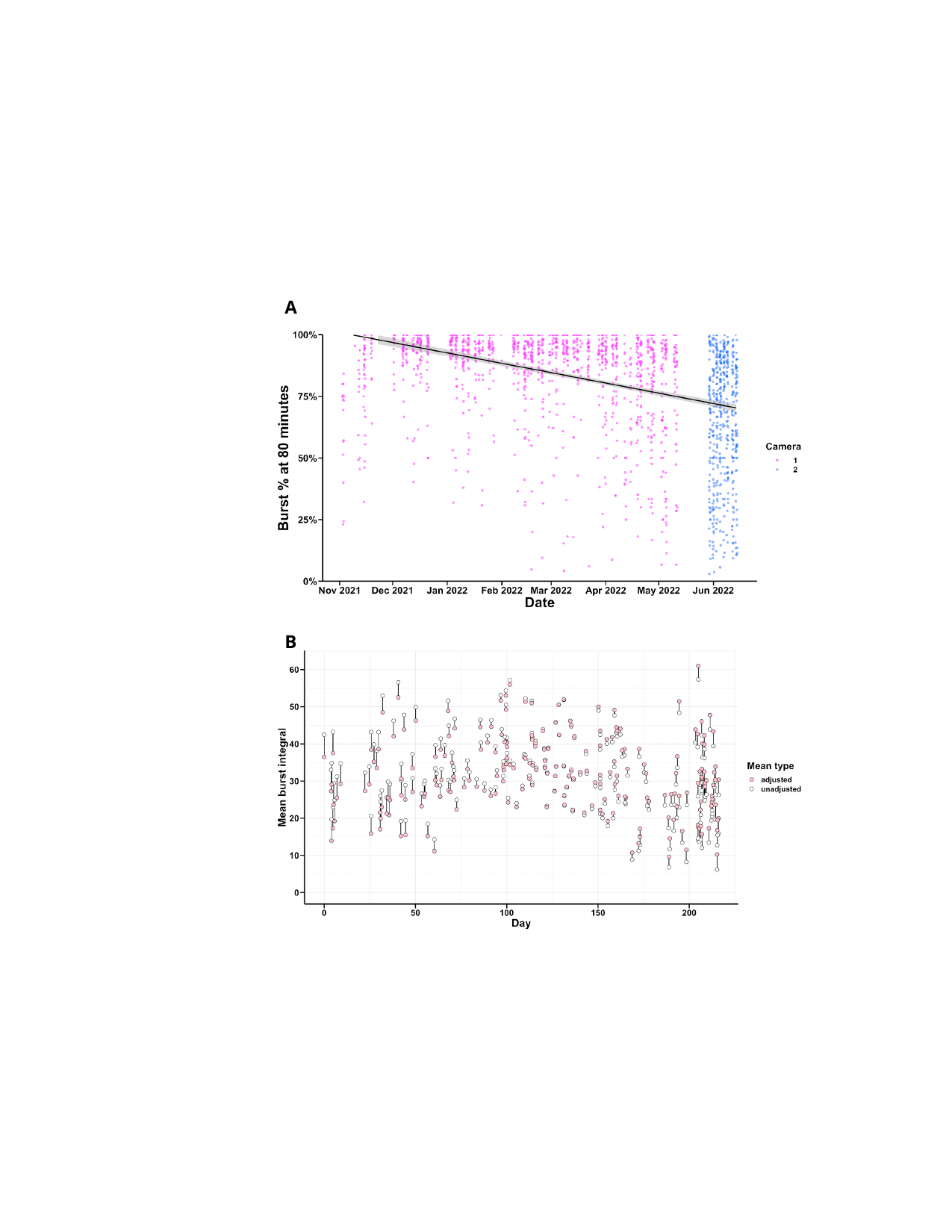

A
B

### Supplemental Figure S4

#### Slide 1
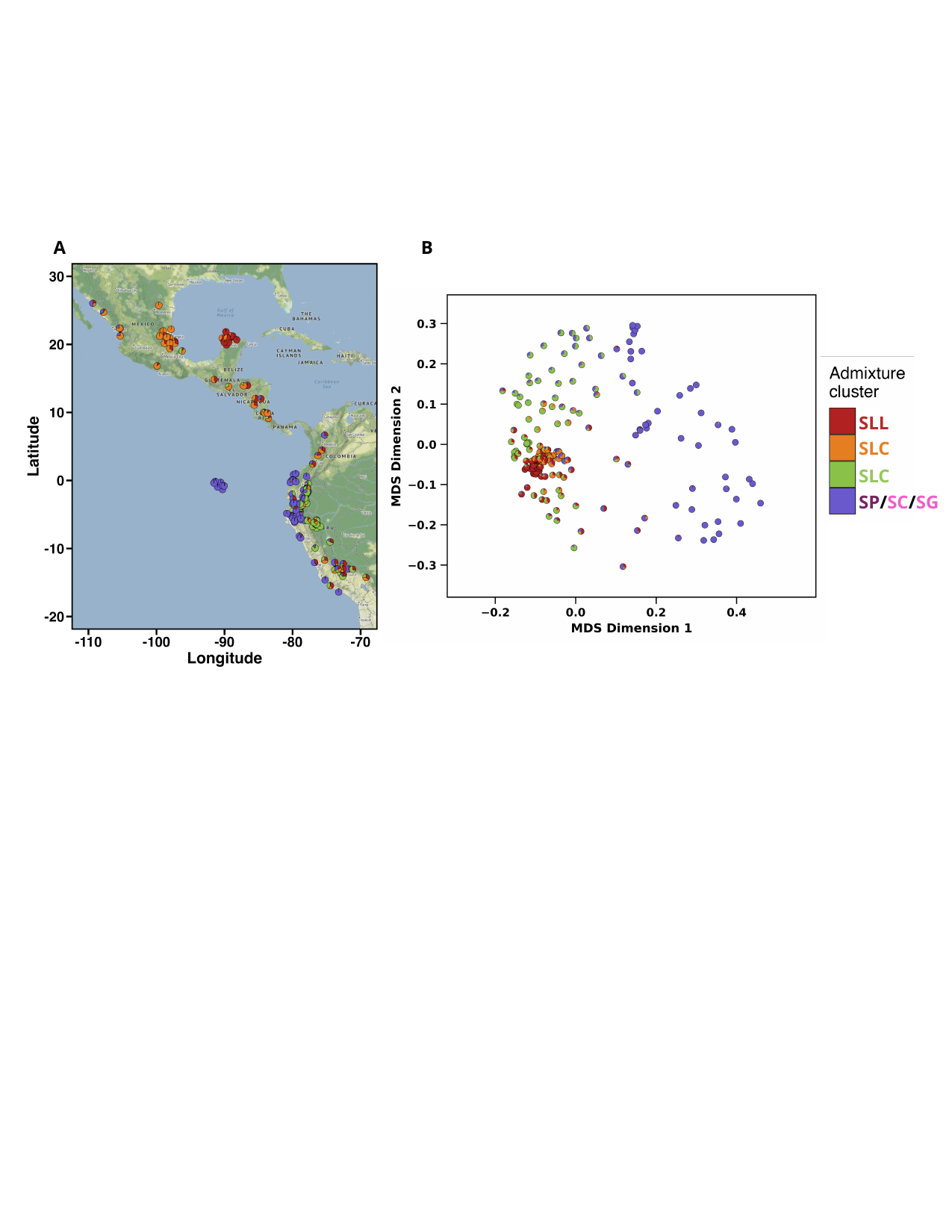

B
A
SLL
SLL
SLC
SLC
SP/SC/SG

### Supplemental Figure S5

#### Slide 1
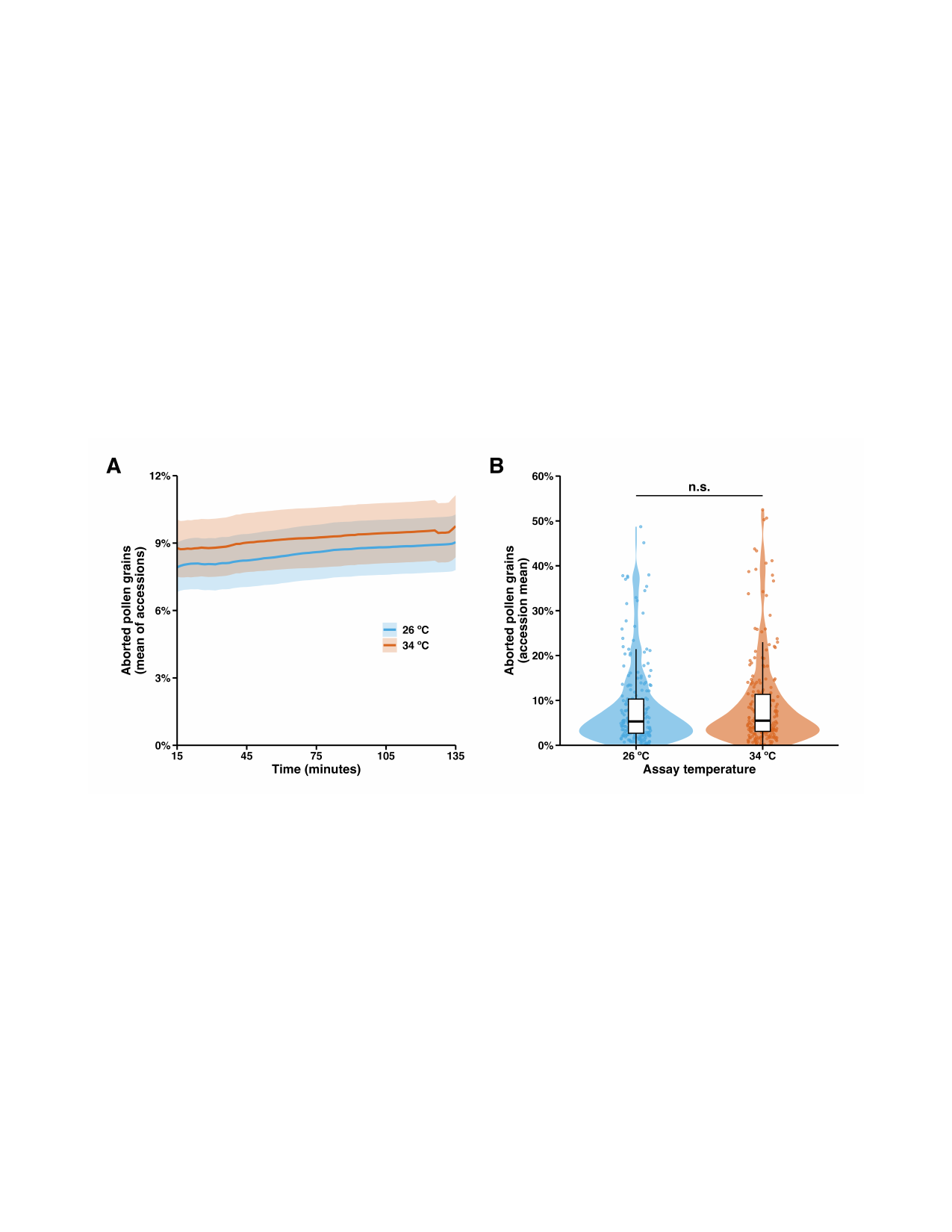

### Supplemental Figure S6

#### Slide 1
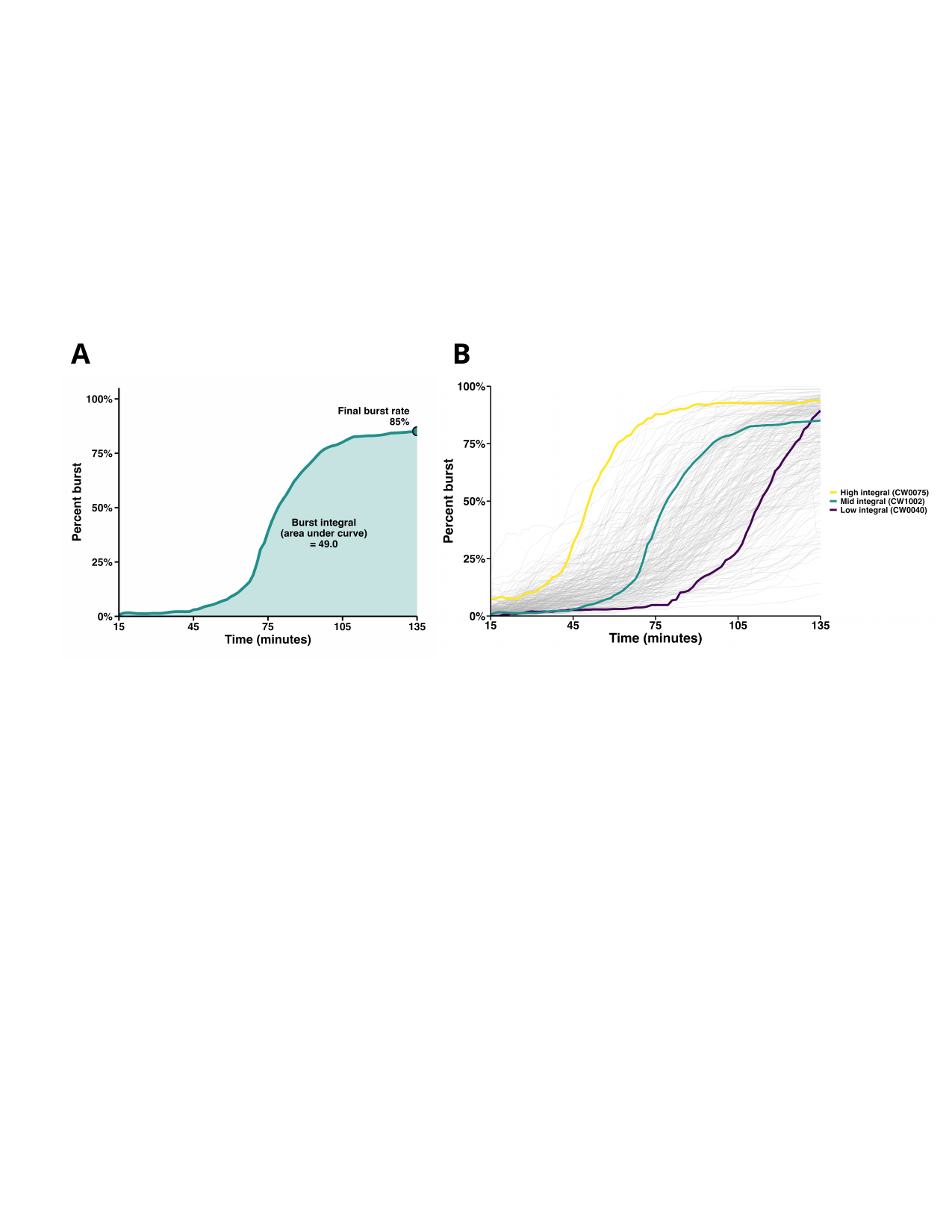

A
B
